# Enteric viruses evoke broad host immune responses resembling bacterial microbiome

**DOI:** 10.1101/2020.10.20.347286

**Authors:** Dallari Simone, Heaney Thomas, Rosas-Villegas Adriana, Jessica A. Neil, Wong Serre-Yu, Judy J. Brown, Urbanek Kelly, Terence S. Dermody, Cadwell Ken

## Abstract

Contributions of the viral component of the microbiome, the virome, to the development of innate and adaptive immunity are largely unknown. Here, we systematically defined the host response in mice to a panel of eukaryotic enteric viruses representing six different families. Most of these viruses asymptomatically infected the mice, the magnitude and duration of which was dependent on the microbiota. Flow cytometric and transcriptional profiling of mice mono-associated with these viruses unveiled general adaptations by the host, such as lymphocyte differentiation and IL-22 signatures in the intestine as well as numerous viral strain-specific responses that persist. Comparison with a dataset derived from analogous bacterial mono-association mice identified bacterial species that evoke an immune response comparable to the viruses we examined. These results expand an understanding of the immune space occupied by the enteric virome and underscore the importance of viral exposure events.

## INTRODUCTION

Our symbiotic relationship with the gut microbiota exemplifies host-microbe coadaptation. In addition to the mutually beneficial exchange of nutrients, intestinal colonization by bacteria shapes the development and function of the mammalian immune system (Honda and Littman, 2012; Round and Mazmanian, 2009). A variety of bacteria evoke context-specific responses that influence the gene expression program of the parenchyma and differentiation of leukocyte subsets (Atarashi et al., 2011; Ivanov et al., 2009; Mazmanian et al., 2005). The outcome of these reactions can be advantageous, as in colonization-resistance to pathogens, or adverse, as in chronic disorders such as inflammatory bowel disease (IBD). Investigations of the host response to individual bacterial species using gnotobiotic animals have led to important insights into the range of immune processes that are fine-tuned by the gut microbiota (Geva-Zatorsky et al., 2017; Sefik et al., 2015a; Tan et al., 2016).

Compared with bacteria, the consequences of intestinal colonization by fungi, protozoans, and viruses on the mucosal immune system are less characterized. Eukaryotic viruses occupy a potentially unique immunologic niche. Viruses, by replicating within mammalian cells, alter signaling cascades and membrane-trafficking pathways, are recognized by nucleic acid sensors and the antigen presentation machinery, and often disseminate to other sites as intracellular passengers. Enteric eukaryotic viruses are detected in healthy infant fecal specimens as early as a few days after birth and become increasingly prevalent and diverse during development (Liang et al., 2020; Lim et al., 2015). Metagenomics analyses of the viral microbiome (virome) have linked various viruses to intestinal disorders such as IBD (Norman et al., 2015; Nyström et al., 2013; Ungaro et al., 2019). Additionally, both transient and persistent infections precede autoimmunity, as observed with the prolonged presence of enterovirus and the development of type 1 diabetes (T1D) (Vehik et al., 2019; Zhao et al., 2017), suggesting viral exposure has long-term immune consequences.

Our studies with murine norovirus (MNV) indicate that eukaryotic viruses can establish a symbiotic relationship with the host akin to commensal bacteria. Germ-free (GF) or antibiotic-treated mice display numerous intestinal defects, including reduced numbers of resident T cells and susceptibility to chemical injury (Round and Mazmanian, 2009). Inoculation with the persistent MNV strain, CR6, reverses these defects by inducing type I interferon (IFN-I), indicating that an antiviral response can provide developmental cues similar to those attributed to the bacterial microbiota (Kernbauer et al., 2014). Furthermore, colonization by MNV is protective in models of childhood enteric bacterial infections and hospital-acquired opportunistic infections (Abt et al., 2016; Neil et al., 2019). Like symbiotic bacteria, MNV triggers adverse outcomes when introduced into a susceptible background. Th1 cytokines induced by MNV cause disease in animal models of IBD (Basic et al., 2014; Bolsega et al., 2019; Cadwell et al., 2010; Matsuzawa-Ishimoto et al., 2017), and the inflammatory gene expression induced by MNV exacerbates bacterial sepsis (Kim et al., 2011). Similarly, MNV and orthoreovirus strain type 1 Lang (T1L), which causes asymptomatic or mild gastrointestinal infection in humans, induces a Th1 response that triggers the loss of immunologic tolerance to dietary gluten in a mouse model of celiac disease (Bouziat et al., 2017, 2018). Rhesus rotavirus (RRV) accelerates autoimmunity in non-obese diabetic mice following recognition by plasmacytoid dendritic cells (pDCs) (Drescher et al., 2015; Pane and Coulson, 2015). These observations may explain the epidemiological association between related viruses and disease in humans (Axelrad et al., 2018, 2019; Bouziat et al., 2017; Pane and Coulson, 2015).

Despite evidence that eukaryotic viruses in the gut have both beneficial and detrimental effects on the host by influencing immune development, a broader characterization of the immune effects of viral exposure is lacking. Administration of antiviral drugs to conventional mice reduces intraepithelial lymphocyte numbers, cytokine levels, and resilience to intestinal injury through IFN-dependent and -independent mechanisms, suggesting that enteric viruses provide a broad range of homeostatic cues to the host (Broggi et al., 2017; Liu et al., 2019; Yang et al., 2016). However, the contribution of individual viruses is unclear.

Here, we conducted an exhaustive cross-comparison of the host response and colonization dynamics of representative enteric viruses. Almost all the viruses we examined evoked a host response in the absence of disease manifestations, and many displayed enhanced capacity to persist in GF mice. Mono-association experiments revealed long-lasting and specific effects of individual viruses on immune cell populations and gene expression. Comparisons with bacteria-associated mice and studies defining the host response to individual bacterial species revealed overlapping yet distinct consequences of viral exposure. These results provide an overview of the immune space occupied by the enteric virome and highlight the wide range of responses that can occur following asymptomatic viral infection.

## RESULTS

### Colonization and bacterial dependence of enteric viruses following a natural route of inoculation

Studies of viral commensalism are hampered by the lack of established animal models. Established models often involve peritoneal or intravenous inoculation of the virus to circumvent local defenses or employ inhibition of antiviral pathways using knockout mice. Another challenge comes from the capacity of segmented filamentous bacterium (SFB) and murine astrovirus, both of which are widespread in institutional vivaria, to inhibit infections by at least some viruses in the intestine (Ingle et al., 2019; Shi et al., 2019). As such, bacterial or viral microbiota may have prevented investigation of certain viruses. These concerns motivated us to perform a side-by-side comparison of viral burden following oral inoculation of conventional, specific-pathogen-free (SPF) and GF mice with different enteric viruses.

We selected a panel of 10 enteric viral strains encompassing six families comprising Groups I, II, III, and IV of the Baltimore classification: two adenoviruses (MAdV1 and 2), an astrovirus (MuAstV), two caliciviruses (MNV CR6 and CW3), a picornavirus (CVB3), two parvoviruses (MVMi and MVMp), and two reoviruses (T1L and RRV). These viruses infect mice, but a detailed time course of infection and corresponding immune response in wild-type C57BL/6 mice following oral inoculation has not been defined for most. Conventional and GF mice inoculated with each virus were monitored for signs of disease and virus in the stool and blood over a 2-month period. We could not recover infectious particles from MNV CW3 at the peak of infection and found that the contents of stool inhibited detection of infectious viral particles, which prevented the use of plaque assays in all conditions (Fig. S1A). A related concern is that detection of infectious particles may be prone to false-negative results once neutralizing antibodies are produced, especially for blood samples. Therefore, we used qPCR, which is a sensitive assay to monitor viral clearance and facilitate comparisons between viruses. T1L and RRV were exceptions for which we used plaque assays, as the multi-segmented nature of the Reoviridae genome confounds quantification by qPCR.

Evidence of disease symptoms, such as diarrhea and hunched posture, were absent in almost all mice, and evaluation of intestinal tissues harvested 28 days post-inoculation (dpi) did not yield evidence of histological abnormalities (Fig. S1B-C). Mice inoculated with CVB3 were the only animals that consistently displayed disease. Despite administering the lowest dose of virus required for seroconversion, ~ 50% of conventional and GF mice did not survive (Fig. S1D). Considering our focus on commensalism, we excluded CVB3 from subsequent studies. We detected replication of each of the remaining nine viruses in both conventional and GF mice (Fig. 1A-B). Although we were unable to detect RRV in stool or blood, we detected anti-RRV neutralizing antibodies, indicating infection (Fig. 1C). MAdV1, MuAstV, and MVMi genomes were detected in the blood at two or more timepoints. Generally, the presence of these viruses in blood predicted their long-term detection in stool (30 dpi).

**Figure 1.**
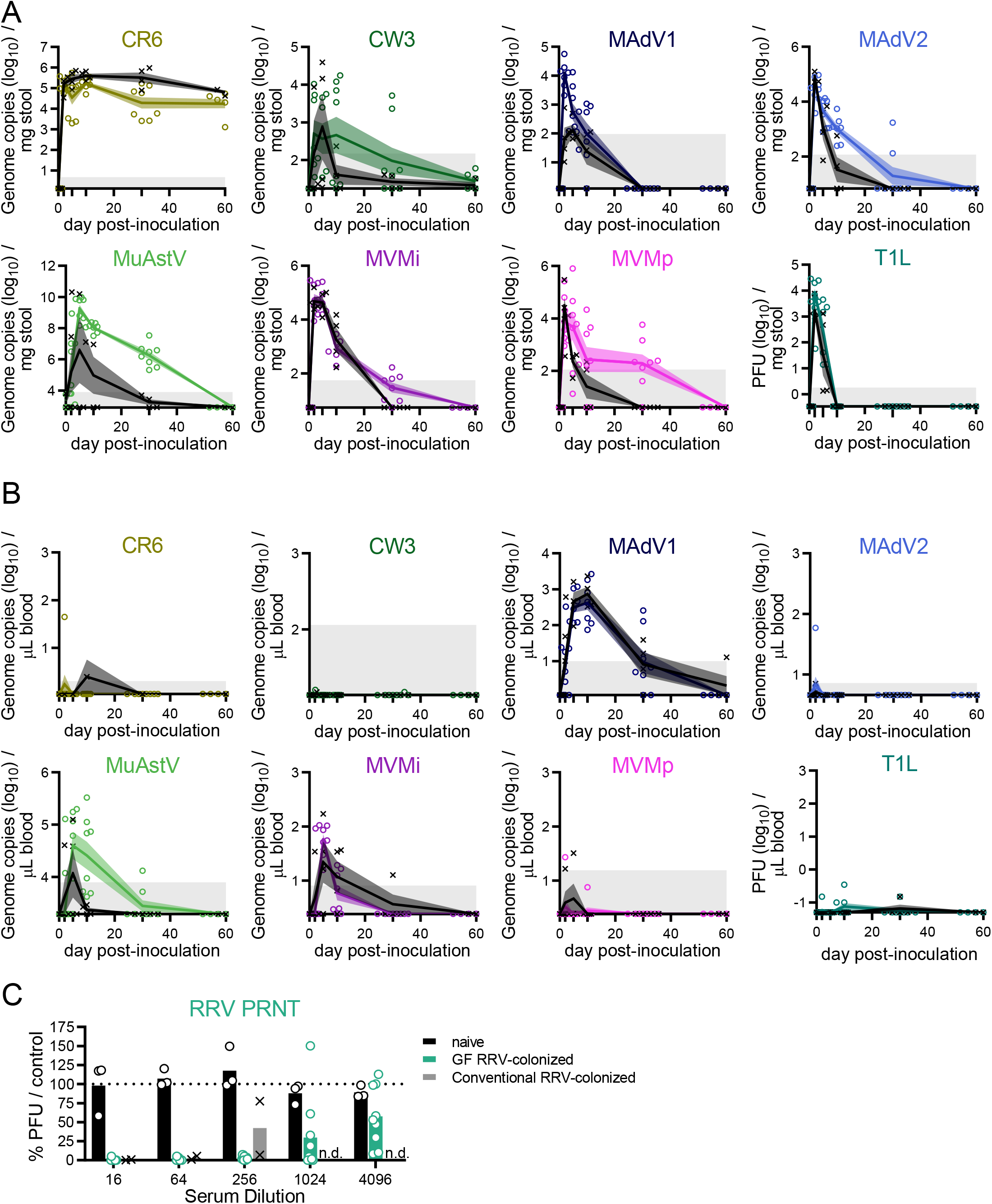
Colonization and Bacterial Dependence of Enteric Viruses Following the Natural Route of Infection. (A-B) Stool (A) and blood (B) were collected at the timepoints shown from conventional (black) and GF mice (colored) inoculated with the virus shown. Viral titers were quantified by plaque assay or qPCR. Symbols indicate individual samples. Lines pass through the mean at each timepoint. Shadowed areas indicate the SEM. Gray areas indicate the limit of detection. N = 4-8 mice per condition, combined from two independent experiments. (C) Neutralizing antibodies in the sera of mice 28 days post-inoculation (dpi) inoculated with RRV were quantified by a plaque-reduction neutralization assay. Reduction in plaque-forming units (PFU) is shown as percent relative to control sera from naïve conventional mice. Results are from 3-9 mice from three independent experiments. n.d.: not determined.

Observations made with antibiotic-treated and GF mice indicate the microbiota is required for optimal infection and transmission by certain enteric viruses (Baldridge et al., 2015; Kane et al., 2011; Kernbauer et al., 2014; Kuss et al., 2011), which we confirmed for MNV CR6. Surprisingly, most of the other viruses displayed similar or enhanced colonization of GF mice, including the closely related MNV CW3 (Fig. 1A-B and S1E). This apparent contradiction can be explained by a recent study showing that bacterial depletion inhibits MNV CW3 infection in one region of the intestine while promoting viral replication in another (Grau et al., 2020). It also is possible that GF mice are more susceptible to viruses because some aminoglycosides used as antibiotics to deplete bacteria from mice elicit an antiviral IFN-I response (Gopinath et al., 2018). MadV1 and T1L reached higher peak titers in GF mice, but the microbiota did not affect the time to clearance (Fig. S1E). In contrast, MNV CW3, MAdV2, MVMi, and MVMp produced similar peak titers but prolonged viral shedding in the stool (Fig. S1E). MuAstV was not uniformly detectable in the stool of conventional mice, perhaps reflecting pre-existing immunity (Yokoyama et al., 2012), but consistently high levels of viral RNA were recovered from GF mice (Fig. 1A-B). Collectively, these data show that exposure to enteric viruses can occur in the absence of overt disease, and many of the viruses chosen for study displayed improved colonization in GF mice. These results, summarized in Table 1, were used to design and interpret the subsequent analysis of the immune response evoked by these viruses.

**Table 1:**
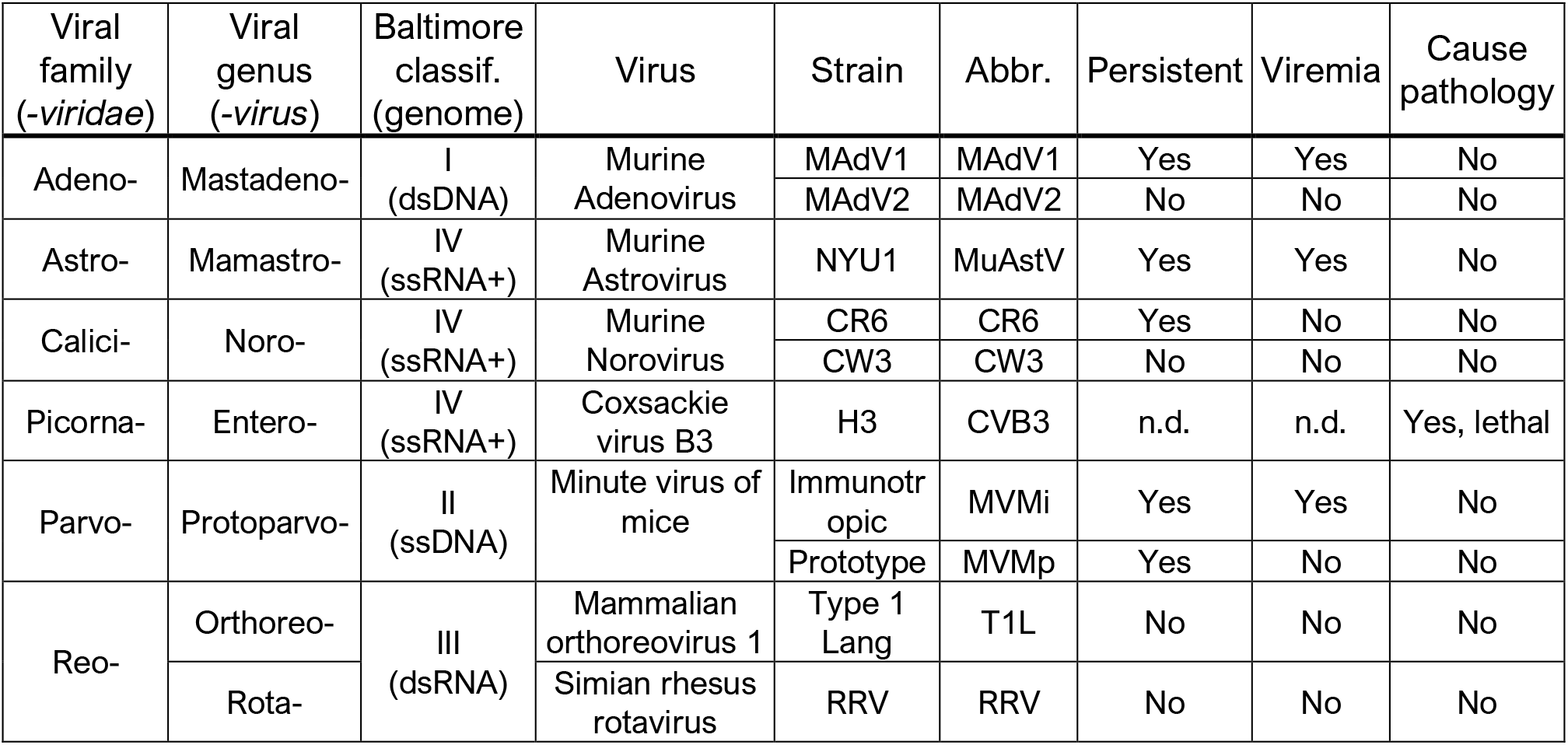
Summary of characteristics of viruses. Taxonomic classification at the family and genus level, Baltimore classification, virus strain names with their abbreviations (Abbr.), and summary of results following inoculation of germ-free (GF) mice from Fig. 1 and S1 for each virus used in this study. Viruses were categorized as persistent if viral nucleic acid was detected at 30 days post inoculation (dpi) in blood or stool following oral inoculation. Viremia is defined as the presence of viral nucleic acid in the blood in at least one time point. The ability to cause pathology is based on the appearance of histological or macroscopic signs of disease, such as lethality or diarrhea. n.d.: not determined.

| Viral family<br>(-viridae) | Viral genus<br>(-virus) | Baltimore<br>classif.<br>(genome) | Virus | Strain | Abbr. | Persistent | Viremia | Cause<br>pathology |
| --- | --- | --- | --- | --- | --- | --- | --- | --- |
| Adeno- | Mastadeno- | I<br>(dsDNA) | Murine<br>Adenovirus | MAdV1 | MAdV1 | Yes | Yes | No |
|  |  |  |  | MAdV2 | MAdV2 | No | No | No |
| Astro- | Mamastro- | IV<br>(ssRNA+) | Murine<br>Astrovirus | NYU1 | MuAstV | Yes | Yes | No |
| Calici- | Noro- | IV<br>(ssRNA+) | Murine<br>Norovirus | CR6 | CR6 | Yes | No | No |
|  |  |  |  | CW3 | CW3 | No | No | No |
| Picorna- | Entero- | IV<br>(ssRNA+) | Coxsackie<br>virus B3 | H3 | CVB3 | n.d. | n.d. | Yes, lethal |
| Parvo- | Protoparvo- | II<br>(ssDNA) | Minute virus of<br>mice | Immunotr<br>opic | MVMi | Yes | Yes | No |
|  |  |  |  | Prototype | MVMp | Yes | No | No |
| Reo- | Orthoreo- | III<br>(dsRNA) | Mammalian<br>orthoreovirus 1 | Type 1<br>Lang | T1L | No | No | No |
|  | Rota- |  | Simian rhesus<br>rotavirus | RRV | RRV | No | No | No |

### A reductionist approach to evaluate responses to viral exposure

To determine whether asymptomatic viral infections are associated with sustained immunological changes, we conducted immune-profiling of mice infected with each virus, a reductionist method similar to that used to define the immunomodulatory activity of individual bacterial species (Geva-Zatorsky et al., 2017; Sefik et al., 2015a; Tan et al., 2016). Although single infections may potentially exaggerate the effect of an individual virus, this approach circumvents concerns about redundancy between viruses in our panel and viral and bacterial members of the microbiota.

We inoculated GF mice perorally with each virus and confirmed infection at 5 dpi. At 28 dpi, six intestinal and extra-intestinal tissues were harvested for analyses by multi-color flow cytometry: colonic and small intestinal lamina propria (cLP and siLP), small intestinal intraepithelial leukocytes (IELs), mesenteric lymph nodes (mLNs), spleen, and lungs. Each sample was analyzed for 32 immune cell subsets based on cell-surface markers and transcription factors. Lymphocyte subsets and functionality were further defined by intracellular staining of six effector cytokines (GRANZYME-B, IL-4, IL-10, IL-17a, IL-22, and IFN-γ) (Fig. S2). Whole colon and small intestine homogenates were subjected to RNA sequencing to examine transcriptional responses. These samples were compared with those prepared in parallel from control GF mice and GF mice colonized with a minimal defined flora (MDF) consisting of a consortium of 15 bacterial strains representing the murine gut microbiota (Brugiroux et al., 2016). These experiments resulted in 462 flow cytometry samples, from which we obtained 21,619 individual immunophenotypes, and 127 transcriptomes.

### Enteric viruses promote changes in immune cell populations

The corresponding fold changes in immune cell populations relative to GF status are shown in Table S1 and the heatmaps in Figure 2A (cLP and siLP) and Figure S3 (IELs, mLNs, lung, and spleen). Viral infection promoted the expansion or contraction of multiple populations, especially in the cLP and siLP. Although each virus had a unique effect, common population changes were altered in a unidirectional manner; we rarely observed a population that increased with one virus and decreased with another. Viruses were observed to modulate as many immune subsets as MDF bacterial microbiota control (Fig. 2A), suggesting that viruses shape intestinal immune responses.

**Figure 2.**
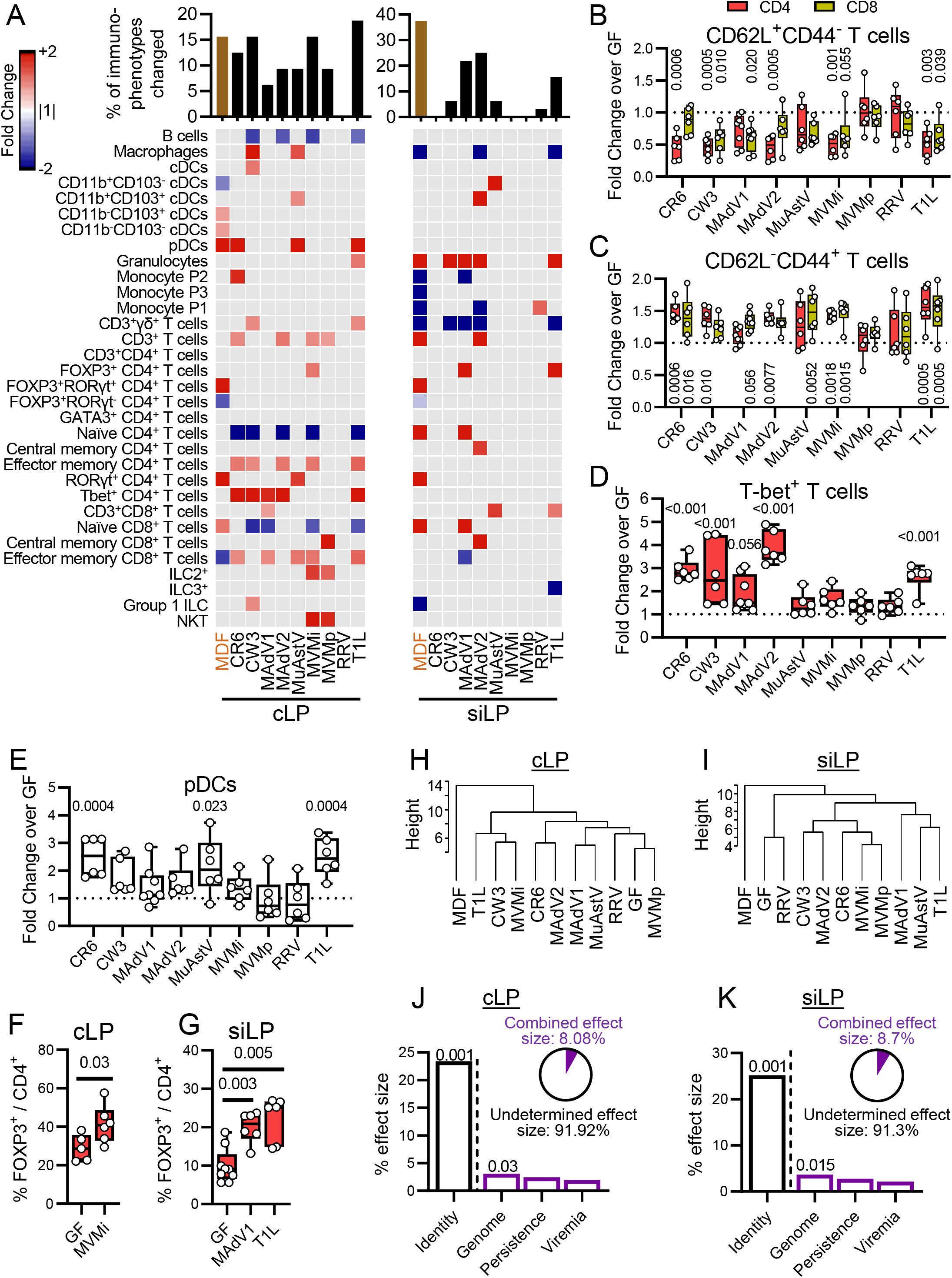
Enteric Viruses Promote Changes in Immune Cell Populations. (A) Heatmap showing average fold-change for cLP and siLP immune populations identified by flow cytometry (using the gating strategy in Fig. S2) for mice inoculated with individual viruses or MDF relative to GF controls with an FDR<0.1. Gray: FDR>0.1. Bar graph on top represents the proportion of immune populations with an FDR<0.1 and a fold change>1.5. (B-E) Fold changes of CD62L^+^CD44^−^ (B), CD62L^−^CD44^+^ (C), T-bet^+^ (D), and pDCs (E) in the cLP CD4^+^ (B-D), CD8^+^ (B-C), or CD45^+^ (E) populations. Each dot represents a single sample. (F-G) Percentage of Foxp3^+^ cells in the cLP (F) or siLP (G) CD4^+^ populations. Each dot represents a single sample. (H-I) Hierarchical clustering of the different conditions based on cLP (H) and siLP (I) population frequencies. (J-K) Effect size determined by db-RDA of viral characteristics: *identity*, *genome*, *persistence*, and *viremia* as explanatory variables of the cLP (J) and siLP (K) population frequency variance. Pie charts represent the combined effect size of *genome*, *persistence*, and *viremia*. Statistical significance was calculated by one-way ANOVA followed by Dunn’s post-hoc analysis and corrected for multiple testing by the Benjamini-Hochberg procedure (A-E) or by non-parametric Mann-Whitney test (F-G).

Our results confirmed several anticipated outcomes, substantiating the validity of our approach. We observed a decrease in CD4^+^ and CD8^+^ naïve T cells (CD62L^+^CD44^−^) and a corresponding increase in CD4^+^ and CD8^+^ effector memory T cells (CD62L^−^CD44^+^) (Fig. 2B-C). We also detected an increase in T-bet^+^ T cells, indicative of a Th1 response (Fig. 2D) (Szabo et al., 2000). Our screen highlighted an increase of macrophages in the cLP in response to MNV CW3, consistent with effects in conventional mice inoculated with this virus (Winkle et al., 2018). Furthermore, at least three enteric viruses induced an expansion of colonic pDCs (Fig. 2E), a population strongly modulated by the bacterial microbiota (Geva-Zatorsky et al., 2017). Despite this commonality, the overlap between mice inoculated with viruses and MDF was limited. One of the most prominent effects of MDF was the induction of FOXP3^−^RORγt^+^ CD4^+^ Th17 cells, but the effect of viral exposure on this population was negligible (Fig 2A). Instead, we observed an increase of FOXP3^+^ CD4^+^ T cells (Tregs) by MVMi in the cLP (Fig. 2F) and by T1L and MAdV1 in the siLP (Fig. 2G). The absence of RORγt within this population (Fig. 2A) suggests that these Tregs are distinct from bacterial-induced peripheral Tregs (Sefik et al., 2015b).

We used hierarchical clustering to define the relative similarity of the overall immune cell composition between conditions (Fig. 2H-I). In both cLP and siLP, MDF was in a clade distinct from individual viruses and the GF control. Viruses did not segregate based on taxonomical relationships, suggesting the immunomodulatory properties observed were marginally intrinsic to a viral family or genus. To quantify how virus-associated variables can explain the variance observed between samples, we conducted a distance-based redundancy analysis (db-RDA) based on shared characteristics (Table 1): genome type (DNA versus RNA), the capacity to persist in the host, defined as detectable virus 30 dpi in blood or stool (*persistence*), and detectable virus in blood (*viremia*). We included the identity of the virus (*identity*) as a benchmark variable in this analysis. Indeed, *identity* was the major explanatory variable, which alone accounted for almost 25% of the variance, supporting the conclusion that individual viruses promote substantially distinct immunomodulatory outcomes (Fig. 2J-K). The second strongest explanatory variable was *genome* type, although the effect size was modest. The combined effect size of the *genome*, *persistence*, and *viremia* variables left much of the variance unexplained, indicating that differences in immune responses to these viruses are likely due to complex interactions between each virus and the host.

Among the other tissue compartments examined, mLNs and lungs displayed the greatest changes in immune cells following viral infection (Fig. S3). Like the intestinal lamina propria, we observed, to a lesser extent, a decrease in naïve and an increase in effector memory CD4^+^ and CD8^+^ T cells in the lungs. Together, these data indicate that enteric viruses influence the immune cell composition of a naïve host, much of which is virus strain-specific. Even for non-persistent viruses, alterations in immune cell frequencies were observed in mice long after the last time point in which viral nucleic acid was detectable.

### Enteric viruses increase cytokine production by immune cells

In parallel with the above analyses, we assessed cytokine production following PMA-ionomycin stimulation of single cell suspensions from each tissue (Fig. 3A, S4A, and Table S1). Inoculation with several viruses led to an increase in cLP T cells producing the Th1 cytokine, IFN-γ (Fig. 3B), which correlated with the increase in T-bet^+^ lymphocytes (Fig. S4B). The increase in IL-17^+^ CD4^+^ T cells was specific to mice colonized with MDF (Fig. 3A). IL-22 is a tissue regenerative cytokine that mediates the protective effect of MNV in models of intestinal injury and bacterial infection (Abt et al., 2016; Neil et al., 2019). Most viruses enhanced IL-22 production by a variety of cLP and siLP lymphoid cells including CD4^+^ T cells, γδ^+^ T cells, and ILCs (Fig. 3A). Quantification of total IL-22^+^ cells using an inclusive CD45^+^ gate in our profiling protocol indicated that cLP infected by six of the viruses and siLP infected by five of the viruses increased the total proportion of IL-22-producing cells (Fig. 3C-D). This IL-22 production by CD45^+^ cells correlated with the proportion of granulocytes and mononuclear phagocytes in the cLP (Fig. S4C). The increase in IL-22^+^ cells was evident in mice that were inoculated with non-persistent viruses, most notably T1L, indicating that alterations in the function of immune cells can be sustained long after the virus is below the threshold of detection (Fig. 3A).

**Figure 3.**
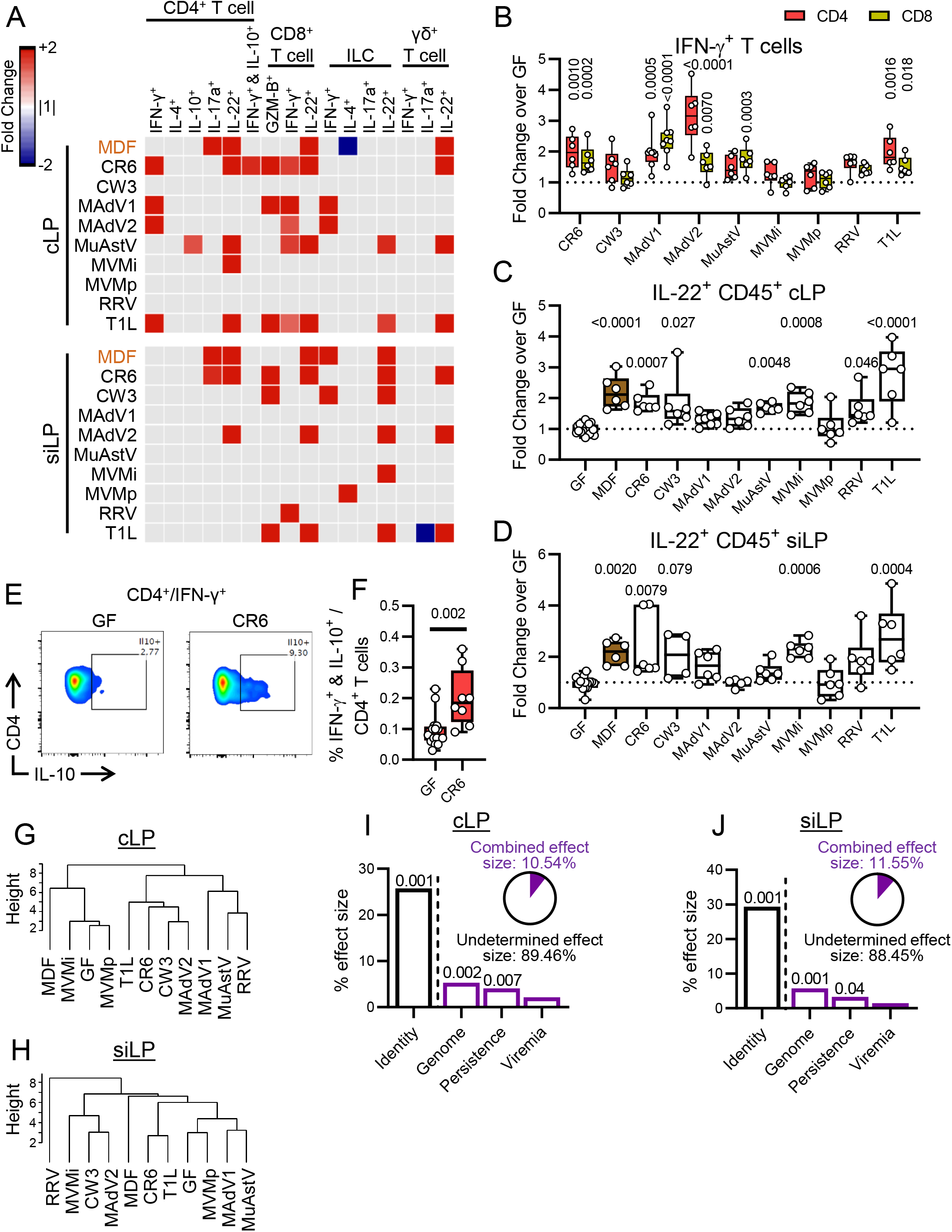
Enteric Viruses Increase Cytokine Production by Immune Cells. (A) Heatmaps showing average fold-change for cytokine-producing immune cells in cLP and siLP identified by flow cytometry for mice inoculated with the viruses shown or MDF relative to GF controls with an FDR<0.1. Gray: FDR>0.1. (B-D) Fold-changes of IFN-γ^+^ (B) and IL-22^+^ (C-D) cells in the cLP CD4^+^ and CD8^+^ (B), cLP CD45^+^ (C), and siLP CD45^+^ (D). (E-F) Representative dot plot (E) and percentage of IFN-γ^+^IL-10^+^ cells in the cLP CD4^+^ T cell population (F). (G-H) Hierarchical clustering of the different microbial associations based on the cLP (G) and siLP (H) cytokine production frequencies. (I-J) Effect size determined by db-RDA of virus as explanatory variables of the cLP (J) and siLP (K) cytokine-producing immune cell frequency variance. Pie charts represent the combined effect size of *genome*, *persistence*, and *viremia*. Statistical significance was calculated by one-way ANOVA followed by Dunn’s post-hoc analysis and corrected for multiple testing by the Benjamini-Hochberg procedure (A-B), by Kruskal-Wallis test followed by Dunn’s post-hoc analysis (C-D), or by non-parametric Mann-Whitney test (F).

Although we did not observe common changes in the capacity to produce cytokines in other tissue compartments as we observed for IL-22 in the lamina propria, we noted several changes in the proportion of cytokine-producing immune cells that were virus strain-specific (Fig. S4A). As an example, IFN-γ^+^IL-10^+^ CD4^+^ T cells (Tr1 cells), a T-helper subset with regulatory functions (Häringer et al., 2009), was increased in mice infected with persistent MNV strain CR6 in cLP and mLN (Fig. 3E-F).

Hierarchical clustering of cytokine production in cLP and siLP cells showed that virus-infected mice did not form clades independent of GF and MDF mice as obviously as they did when analyzing immune cell populations based on cell-surface markers and transcription factors (Fig. 3G-H). As with the prior analyses, viruses from the same families did not uniformly cluster together, and the major explanatory variables for cytokine production were *identity*, followed by *genome* (Fig. 3I-J). Together, these results indicate that virus-infected mice display increases in cytokine-producing immune cells that are both common and virus-strain specific.

### Intestinal transcriptome of virus-infected mice

Of genes profiled in the colon and small intestine, 497 and 355, respectively, displayed differential expression (DE) in at least one virus-infection condition compared with GF mice (≥ 2-fold, p value ≤ 0.01) (Fig. 4A-C, Table S2A-B). In comparison, 146 and 92 genes in the colon and small intestine displayed differential expression in MDF-colonized mice and minimally overlapped with the virus-induced expression changes (Fig. 4D-E, Table S2C-D). Gene ontology (GO) analyses showed that viral infection influenced a wide range of immune-related pathways, especially in the colon (Fig. 4F-G). Viral infection was associated with antiviral immunity pathways, such as *defense response to virus* and *cellular response to interferon-beta*. The enrichment for genes associated with IFN-γ is consistent with the flow cytometry data identifying a Th1 response. Both MDF and viruses were associated with B cell activation and bacterial response pathways. The enrichment of DE genes involved in metabolic processes was specific to MDF, perhaps reflecting the nutrient exchange between host and bacteria.

**Figure 4.**
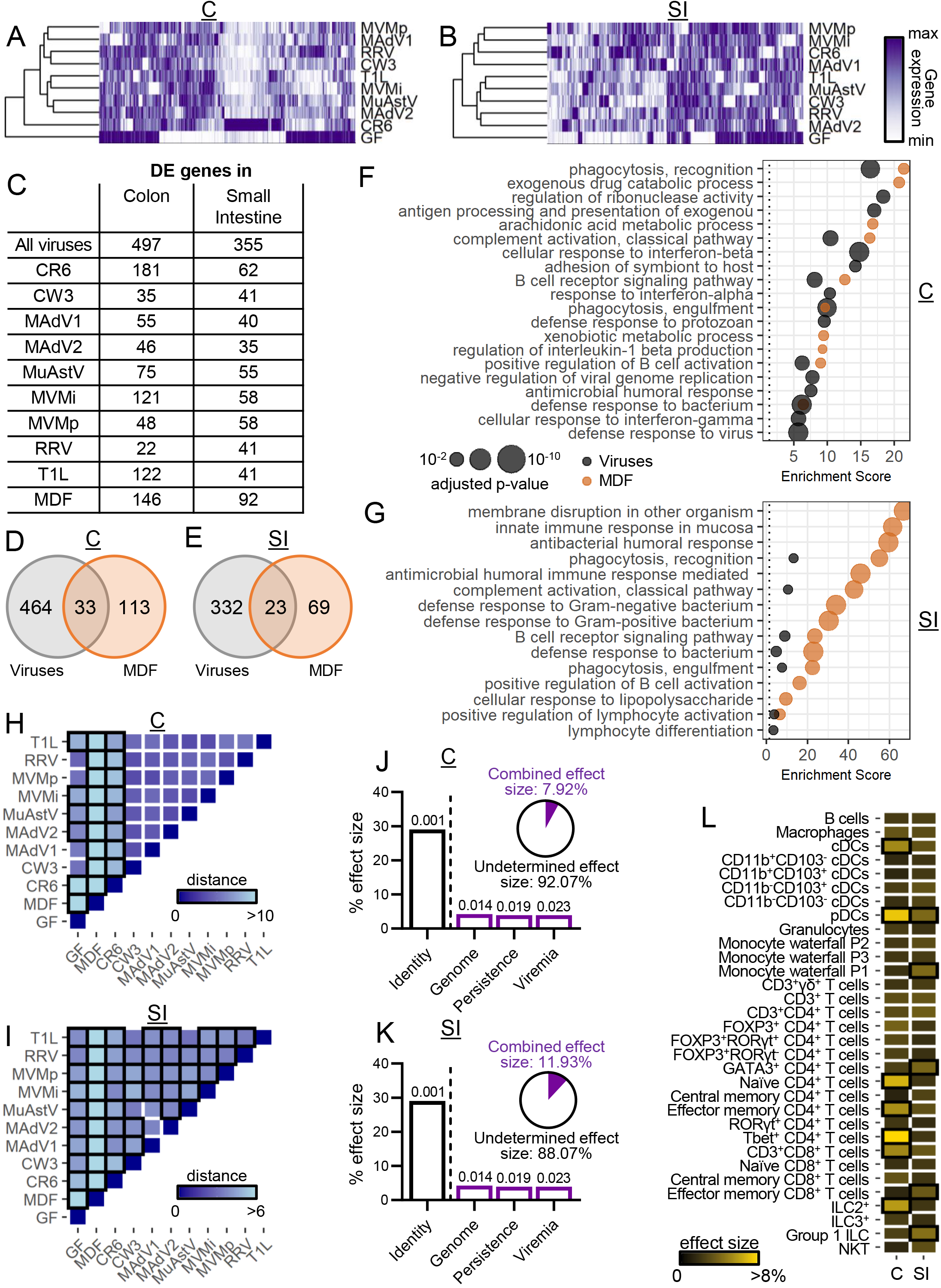
Intestinal Transcriptome of Virus-Infected Mice. (A-B) Heatmaps showing DE genes (|average fold-change over GF|≥2 and unadjusted p-value≤0.01) in the colon (A) and small intestine (B) of virus-infected mice compared with GF mice. C: colon; SI: small intestine. (C) Number of DE genes in the colonic and small intestinal transcriptome for each condition compared with GF mice. (D-E) Venn diagrams depicting the number and overlap of DE genes in all virus-infected and MDF-associated mice in the colon (D) and small intestine (E). (F-G) Top 15 most highly enriched biological process GO terms for the DE genes in the colon (F) and small intestine (G) of virus-infected and MDF-associated mice. (H-I) Heatmaps showing the Euclidean distances between group centroids of DE genes in the colon (H) and small intestine (I) comparing each condition. Boxes outlined in black indicate significant differences (PERMANOVA<0.05). (J-K) Effect size determined by db-RDA of virus characteristics as explanatory variables of the DE gene variance in the colon (J) and small intestine (K). Pie charts represent the combined effect size of *genome*, *persistence*, and *viremia*. (L) Effect size determined by db-RDA of immune population frequencies from Figure 2 on DE gene variance in the colon and small intestine. Boxes outlined in black indicate p-value<0.05.

Permutational multivariate analysis of the variance after principal component analysis (PCA) confirmed that the transcriptional responses to viruses differed significantly from that of GF and MDF conditions and that each virus induced a distinct gene expression pattern (Fig. 4H-I and S5A-B). The major explanatory variables of the variance between samples were again *identity* followed by *genome* (Fig. 4J-K). Because much of the transcriptome variance was unexplained, we determined whether the immune cell composition and cytokine production (described in Figs. 2 and 3) correlated with differences in gene expression between conditions. DC and T cell subsets were major explanatory variables and included cell types with recognized functions in antiviral responses such as Tbet^+^ CD4^+^ T cells and pDCs (Fig. 4L). Among the cytokines tested, only IL-22 was a significant explanatory parameter (Fig. S5C), likely reflecting the role of this cytokine in coordinating antimicrobial gene expression (Keir et al., 2020). Collectively, these results correlate well with our flow cytometry data and reveal responses common to multiple viruses, while also underscoring the importance of investigating the immune effects of individual virus strains, which cannot be predicted based on taxonomic features alone.

### Intestinal gene expression common and specific to individual viruses

We next identified specific genes and pathways associated with each virus individually and those in common. We observed 15 and three differentially regulated genes in the colon and small intestine, respectively, that were shared by at least half of the viruses studied, including immunoglobulin genes *Igha*, *Igkc*, *Iglc1*, *Jchain*, and *Pou2af1* (Fig. 5A-B). This finding is consistent with the increased expression of genes associated with B cell activation (Fig. 4F-G) as well as previous findings that MNV CR6 enhances local and systemic antibody production in GF mice and that IgA production is frequently observed during enteric viral infections (Blutt and Conner, 2013; Kernbauer et al., 2014).

**Figure 5.**
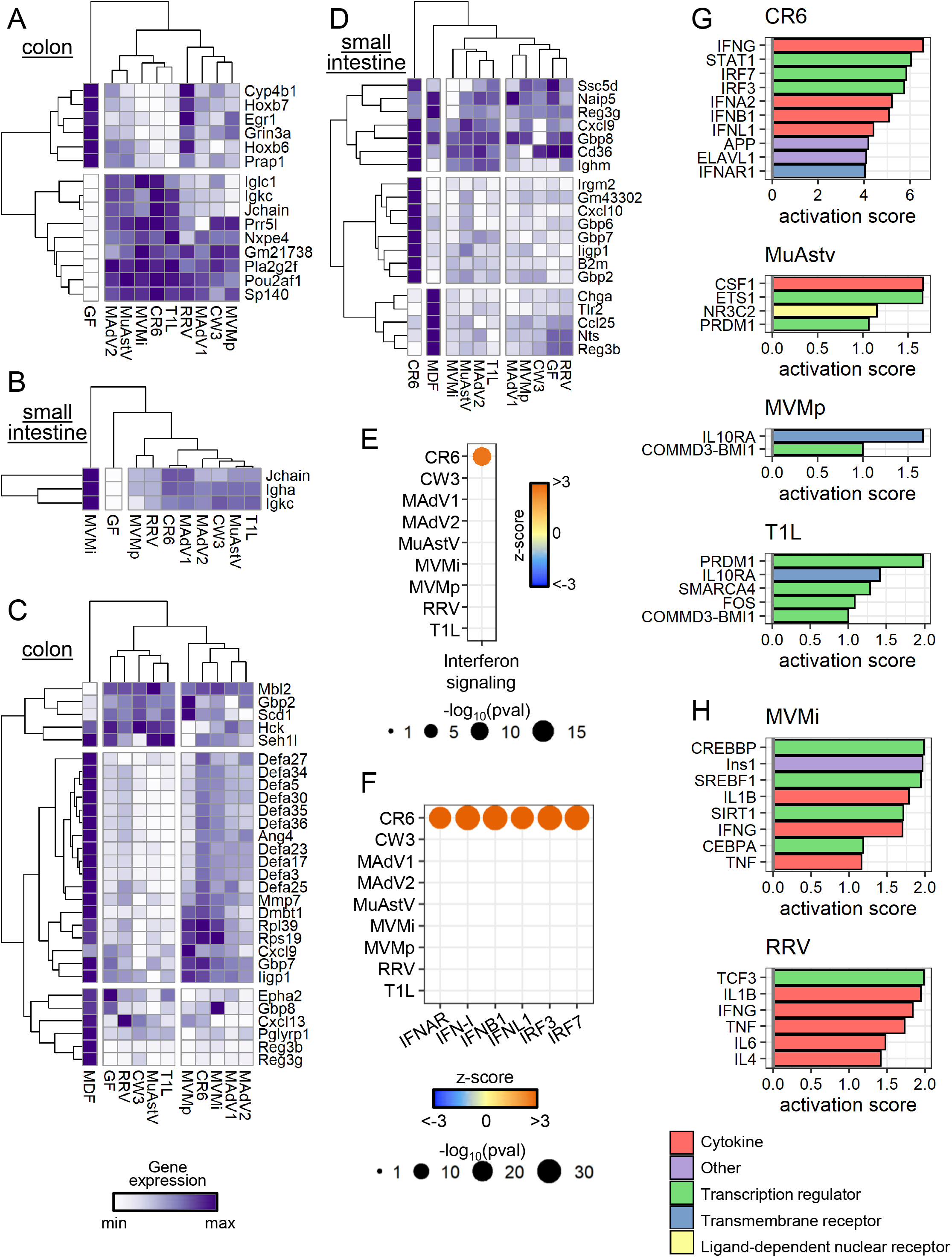
Intestinal Gene Expression Common and Specific to Individual Viruses. (A-B) Heatmaps displaying normalized expression values of DE genes (average fold-change over GF≥2 and unadjusted p-value≤0.01) modulated by at least five viruses in the colon (A) and small intestine (B). (C-D) Heatmaps displaying normalized expression values of DE genes (average fold-change over GF≥1.5 and unadjusted p-value≤0.01) annotated in GO:0050829, GO:0050830, and GO:0061844 in the colon (C) and small intestine (D). (E-F) Ingenuity pathway analysis (IPA) of the colonic transcriptome of virus-infected mice for enrichment of DE genes involved in IFN signaling (E) or key molecules in the IFN pathway (F). (G-H) Colonic (G) and small intestinal (H) DE genes were analyzed by IPA for upstream regulators. Top 10 upstream regulators for each virus with an activation score>1 are depicted.

Increased expression of antimicrobial genes is a hallmark of intestinal colonization by symbiotic bacteria (Geva-Zatorsky et al., 2017; Hooper et al., 2001). We examined expression of antimicrobial genes during viral infection by constructing an antimicrobial gene set in which genes annotated in GO:0050829 (defense response to Gram-negative bacterium), GO:0050830 (defense response to Gram-positive bacterium), and GO:0061844 (antimicrobial humoral immune response mediated by antimicrobial peptide) were pooled. Expression of numerous antimicrobial genes was increased in virus-infected mice compared with GF controls (≥ 1.5-fold change, p ≤ 0.01) (Fig. 5C-D). However, the overall response was not as strong as that induced by MDF. Nonetheless, there were transcripts induced exclusively by viruses, including mannose-binding protein C (*Mbl2*) and the interferon-inducible GTPases, *Iigp1*, *Irgm2*, and *Gbp2*.

MNV CR6 but not MNV CW3 fortifies the intestinal barrier by inducing a local IFN-I response (Kernbauer et al., 2014; Neil et al., 2019). IFN-I (IFN-α and −β) and type III interferon (IFN-λ) are antiviral cytokines produced in response to viral nucleic acid that evoke a similar set of interferon-stimulated genes (ISGs), which we collectively term here as an “IFN signature”. Consistent with our previous findings, colonization with MNV CR6 but not MNV CW3 was associated with an IFN signature (Fig. 5E). Surprisingly, no other virus from our panel yielded an IFN signature, despite high levels of viral nucleic acid produced by some of them, such as MuAstV. Moreover, only MNV CR6 was associated with increased transcription of ISG regulators (Fig. 5F). We confirmed that expression of representative ISGs *Isg15*, *Ifit1*, and *Oas1a* was increased only in mice colonized with MNV CR6 (Fig. S6A).

In contrast to IFN-I, IL-22 should regulate expression of DE genes for multiple viruses based on its effect size on transcriptional variance (Fig. S5C). To test this prediction, we used Gene Set Enrichment Analysis (GSEA) to determine whether transcripts altered in the intestine of *Il-22*^−/−^ mice (Gronke et al., 2019) were differentially regulated in virus-infected mice. This analysis confirmed that most virus-infected mice produced an IL-22 signature. (Fig. S6B-C).

We used Ingenuity Pathway Analysis (IPA) to identify additional regulators in the following categories: *cytokine*, *ligand-dependent nuclear receptor*, *transmembrane receptor*, *transcript regulator*, and *other*. In the colon, four viruses were associated with such regulators. IFN-related factors were the main regulators associated with MNV CR6, whereas MuAstV, MVMp, and T1L upregulated other pathways (Fig. 5G). PRDM1, also known as BLIMP1, is a regulator of terminal B-cell differentiation (Shaffer et al., 2002) and influenced transcriptional responses to MuAstV and T1L, supporting a role for viruses in B cell development. The association of MuAstV and macrophage differentiation factor CSF-1 is consistent with the observation that MuAstV-colonized mice displayed an increase in cLP macrophages (Fig. 2A). In the small intestine, genes induced by pro-inflammatory cytokines such as IL-1β, IFN-γ, and TNF-α were enriched in mice infected with either MVMi or RRV (Fig. 5H). Other factors identified by this approach have been implicated in immunity in some settings. For example, insulin (Ins1), which was associated with MVMi infection, is involved with IFN-γ in a feedback loop to promote the effector CD8^+^ T cell response to murine cytomegalovirus infection (Šestan et al., 2018). Therefore, in addition to the classic ISGs downstream of IFN-I that we observe for MNV CR6, viral exposure induces the expression of genes regulated by a range of signaling molecules and pathways required for mucosal immunity.

### Intestinal transcriptomes of virus-infected mice are enriched for bacterial microbiome gene signatures

We used a GSEA strategy analogous to a previously described approach (Godec et al., 2016) to compare the transcriptome of virus-infected mice with gene-expression signatures of mice monocolonized with 53 individual species of the bacterial microbiota (Geva-Zatorsky et al., 2017) (Fig. 6A-D, Table S3). Colonic transcripts of MDF-colonized mice in our study were positively enriched for genes upregulated in microbiota-replete conditions (conventional SPF mice) (Geva-Zatorsky et al., 2017), indicating concordance in the positive controls (Fig. 6A). Similarly, the small intestinal transcriptome of MDF-colonized mice displayed a negative enrichment score for genes downregulated in conventional SPF mice (Fig. 6D). Twenty of the 53 bacterial species displayed a relationship with one or more viruses using this approach. We identified 91 virus-bacterium pairs, with 60 displaying the same directionality of regulation (i.e., positive enrichment of upregulated bacterial gene sets in virus-associated transcripts and negative enrichment of downregulated bacterial gene sets in virus-associated transcripts) (Fig. 6A-D). Certain bacterial gene sets displayed exclusive pairing with one virus, as observed with several *Bacteroides* and *Parabacteroides* species and T1L in the colon. This consistent pairing between T1L and the Bacteroidales order suggests that this virus induces a similar reaction to colonization by this prototypical group of commensal bacteria. The MNV CR6-induced gene set also was paired with multiple bacterially upregulated gene sets in the colon. We observed a particularly strong enrichment for the *Enterococcus faecalis* signature, a facultative anaerobic bacterium of the Enterococcaceae family (Fig. 6E).

**Figure 6.**
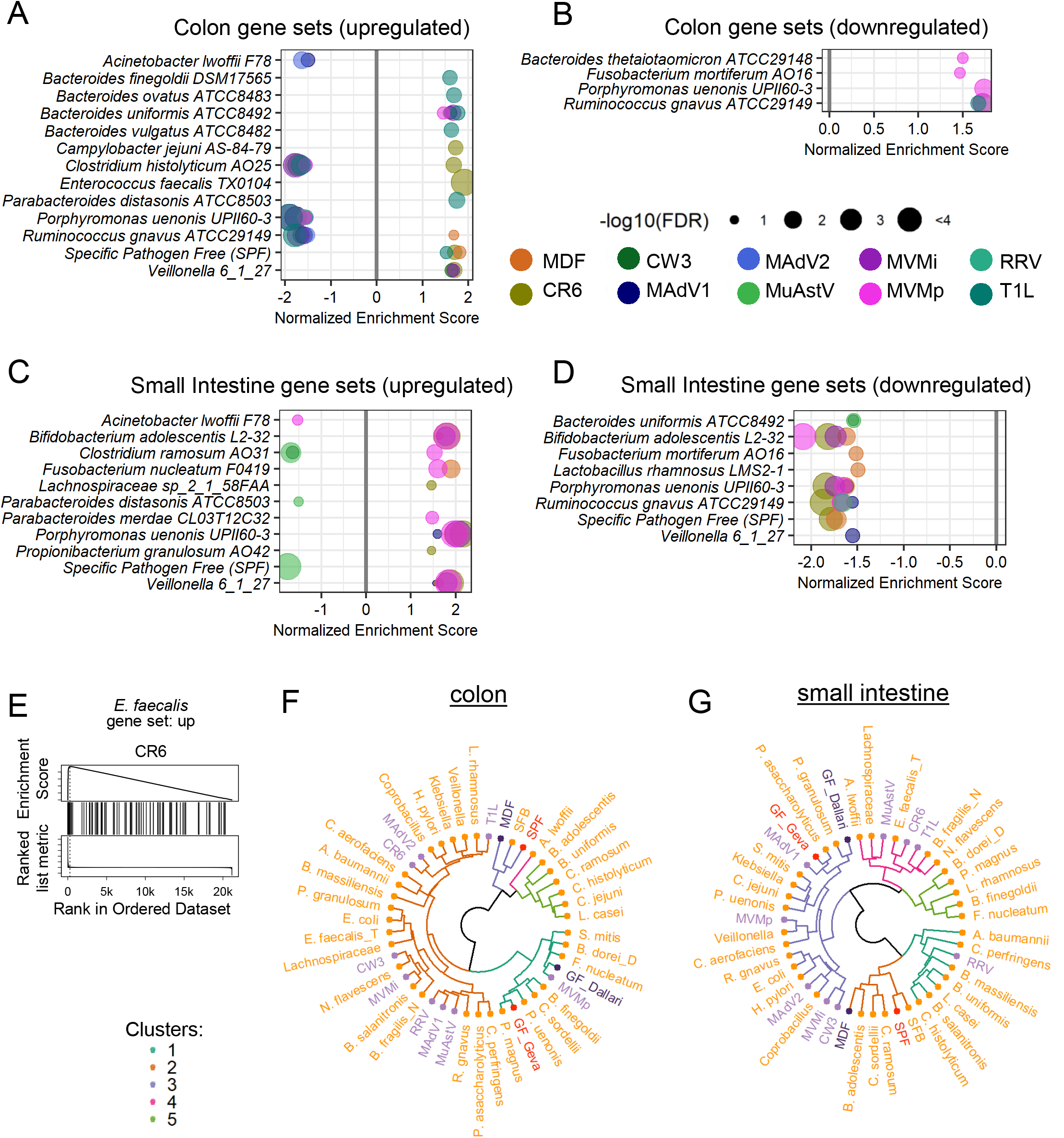
Intestinal Transcriptomes of Virus-Infected Mice Are Enriched for Bacterial Microbiome Gene Signatures. (A-D) Colonic (A-B) and small intestinal (C-D) transcriptomes from virus-infected mice compared with gene expression signatures of bacterially colonized mice by GSEA. Gene sets upregulated following colonization by bacteria are depicted in A and C; downregulated gene sets are depicted in B and D. (E) GSEA plot showing enrichment of the *E. faecalis* upregulated gene set in the colonic transcriptome of mice infected with MNV CR6. (F-G) Hierarchical clustering of the immune population frequencies described in Table S4. Purple: viruses; dark purple: MDF and GF from this study; orange: bacteria; pink: conventional SPF and GF from Geva-Zatorsky et al.

Lastly, we compared our intestinal flow cytometry data with results gathered using mice monocolonized with bacteria (Geva-Zatorsky et al., 2017). Due to differences in gating strategies and markers used to identify cell types, we restricted our comparison to 16 immune cell subsets that were quantified in a similar manner across the datasets (Table S4). Hierarchical clustering using the z-scores of the two datasets indicated that the GF groups from both datasets clustered together, as did MDF from our study and SPF from Geva-Zatorsky et al (Fig. 6F-G). Most bacteria and viruses clustered together with GF or in neighboring clades distinct from MDF and SPF, suggesting the contribution of a specific bacterium or virus, when present alone, accounts for only a modest fraction of the total microbiota-dependent effects on immune cell composition. The viruses were interspersed among bacteria rather than clustered together in a single clade, indicating that virus-induced changes to immune cell frequencies does not reflect a uniform immunological response to viruses distinct from that evoked by bacteria.

## DISCUSSION

In this study, we investigated whether asymptomatic or subclinical infections of the gastrointestinal tract by eukaryotic viruses shape the mucosal immune system, as has been demonstrated for numerous bacterial members of the microbiota (Honda and Littman, 2012; Round and Mazmanian, 2009). Only one of the 10 viruses chosen for study led to illness or death, allowing us to define the immune effects of nine viruses in the absence of disease.

In the process of establishing virus infection models, we made several observations about the dynamics of infection. First, we found that nucleic acid of several viruses remains detectable in stool or blood for a prolonged interval. Unlike retroviruses and herpesviruses, members of these viral families are not known to establish latency. Observations with measles virus infections indicate that viral antigens and RNA can persist, even for viruses that do not establish latency or integrate DNA copies into the host genome (Griffin, 2020). For some viruses, this type of persistence could be mediated by immune evasion, as proposed for MNV (Lee et al., 2019; Tomov et al., 2017). Regardless of the mechanism, we found that the microbiota had a strong effect on persistence. While antibiotic treatment hinders infection by some enteric viruses (Baldridge et al., 2015; Kernbauer et al., 2014; Kuss et al., 2011), our data showed that not all viruses benefit from the presence of bacteria. An important future goal is to determine whether this resistance to infection displayed by conventional mice reflects the presence of specific autochthonous viruses or bacteria in the gut (Ingle et al., 2019; Shi et al., 2019).

Our flow cytometric and transcriptional analyses were well correlated and support the hypothesis that asymptomatic colonization by enteric viruses has consequences for the host. Although each virus was associated with a unique immune profile following oral inoculation of GF mice, there were a few recurrent themes. Viral infection generally promoted the differentiation of lymphocytes, specifically maturation of T cells and Th1 polarization. Laboratory mice display deficiencies in mature T cells due to the absence of exposure to infectious agents while housed in SPF conditions (Beura et al., 2016; Lin et al., 2020; Yeung et al., 2020). In this context, it is notable that wild mice and pet-shop mice, which have a more mature lymphocyte compartment, are seropositive for viruses closely related to those in our panel (adenovirus, MNV, parvovirus, reovirus, and rotavirus) (Beura et al., 2016). These common enteric viruses may contribute to immune maturation in the natural environment.

We find it noteworthy that the immune effects of a given virus could not be explained by qualitative features alone. Closely related viral strains evoked distinct responses in most of the parameters we assessed. The nucleic acid composition of the viral genome (DNA versus RNA) contributed modestly but reproducibly to the variance, whereas viral dissemination and persistence did not appear to explain differences between conditions. Accordingly, one remarkable finding was that changes to immune cells and gene expression patterns were readily observed in mice in which viral nucleic acid was no longer detectable. If this sustained effect of viruses translates to humans, then cross-sectional metagenomics studies of patient cohorts would miss potentially meaningful exposures to viruses that occurred prior to disease onset. Longitudinal virome analyses of children genetically susceptible to T1D identified an inverse relationship between early life adenovirus and circovirus exposure with subsequent appearance of serum autoantibodies (Vehik et al., 2019; Zhao et al., 2017). Thus, we advocate prospective and longitudinal sampling for virome-association studies when possible.

Based on our prior studies with MNV, we anticipated that at least some virus-infected mice would display an IFN signature. Instead, we observed an increase in IL-22-producing cells and an IL-22-mediated gene-expression pattern following infection by several of the viruses in our panel. IL-22 functions in intestinal homeostasis and expression of antimicrobial genes (Gronke et al., 2019; Keir et al., 2020). We think it possible that IL-22 induction offsets damage caused by enteric viruses, thereby facilitating a commensal relationship.

A comparison between our results and an analogous dataset gathered using bacterial monocolonization identified virus-bacterium pairs that stimulate overlapping responses by the host. For example, *E. faecalis* and MNV CR6 shared a colonic gene expression signature, which increased our confidence in the approach because these two infectious agents also share the capacity to confer protection in the DSS model of intestinal injury (Kernbauer et al., 2014; Neil et al., 2019; Takahashi et al., 2019; Wang et al., 2014). Several of the bacteria that evoke an immune response overlapping with viruses are implicated in disease, such as *Ruminococcus gnavus* and *Bacteroides vulgatus* in IBD (Hall et al., 2017; Png et al., 2010; Rath et al., 1999). It will be interesting to test the role of the matching viruses in animal models in which disease is dependent on these bacteria (Bloom et al., 2011; Ramanan et al., 2014, 2016; Yu et al., 2020).

Our survey was restricted to a limited number of viruses and, therefore, we were not able to capture the vast diversity of viruses found in humans. Unlike bacteria isolated from the human gut, which almost always colonize GF mice, many medically important viruses display narrow species tropism or altered virulence when inoculated into mice. A broader survey of viruses will likely identify additional cell types and pathways influenced by viral infection. Another limitation is that we chose a single-infection approach to identify direct responses and avoid missing immune effects that overlap with the existing microbiota. This approach also enabled our *in-silico* comparison of virus-induced immune responses with those induced in mice monocolonized with bacteria.

We envision two situations in which our results can guide studies investigating how the enteric virome modulates immunity in the presence of bacteria. First, mice associated with defined flora can be used to assess the immune effects of individual bacteria within a complex community (Fischbach, 2018). This synthetic ecology approach could incorporate viruses with immunogenic potential from our panel to better reflect the complexity of the real-world microbiome. Second, we advocate testing the role of these and other viruses in animal models of inflammatory diseases, many of which are thought to be dependent on bacterial members of the microbiota. Although the Th1 response to MNV CR6 is inconsequential in wild-type C57BL/6 mice, mutation of IBD-susceptibility gene *ATG16L1* sensitizes the intestinal epithelium to the otherwise subtle effect of viral infection (Cadwell et al., 2010; Matsuzawa-Ishimoto et al., 2017). Observations in studies of MNV-infected mutant mice allowed us to identify homeostatic mechanisms involved in barrier integrity that are conserved in humans (Cadwell et al., 2008; Matsuzawa-Ishimoto et al., 2020). Thus, incorporating viruses into genetic disease models can reveal vital pathways that promote health.

Our findings indicate that eukaryotic viruses in the gut have unappreciated immunomodulatory capacity in addition to well-recognized roles as causative agents of gastroenteritis. The reaction to viral infection could be beneficial in the appropriate setting, as demonstrated by proof-of-principle experiments showing that MNV and MuAstV strains administered prophylactically protect mice from enteropathogenic *E. coli* (Cortez et al., 2020; Neil et al., 2019). There is precedent for manipulation of the gut virome for therapeutic purposes. Oral poliovirus vaccine provides cross-protection against other pathogens, which has been used as a rationale to administer this attenuated virus instead of inactivated vaccine in polio-endemic regions (Upfill-Brown et al., 2017). Our ongoing studies using animal models will enable future safety and efficacy assessments of virome-based interventions.

## ACKNOWLEDGMENTS

We wish to thank Dr. Julie Pfeiffer (UT Southwestern), Dr. Jason G. Smith (University of Washington), Dr. David Pintel (University of Missouri), Dr. Peter Tattersall (Yale University), Dr. Harry B Greenberg (Stanford University), and Dr. Kathy McCoy (University of Calgary) for sharing reagents with us and providing advice regarding culturing techniques. We wish to thank the NYU Grossman School of Medicine Flow Cytometry and Cell Sorting, Microscopy, Genome Technology, and Histology Cores for use of their instruments and technical assistance (supported in part by National Institutes of Health [NIH] grants P31CA016087, S10OD01058, and S10OD018338). We also wish to thank Margie Alva, Juan Carrasquillo, and Beatriz Delgado for technical assistance with gnotobiotics. This research was supported by NIH grants DK093668 (K.C.), AI121244 (K.C.), HL123340 (K.C.), AI130945 (K.C.), R01 AI140754 (K.C.), F31 DK108562 (J.J.B.), T32 HL007751 (J.J.B.), R01 AI038296 (T.S.D.), R01 DK098435 (T.S.D.), and a pilot award from the NYU Cancer Center grant P30CA016087 (K.C.). Additional support was provided by the Faculty Scholar grant from the Howard Hughes Medical Institute (K.C.), Crohn’s & Colitis Foundation (K.C.), Merieux Institute (K.C.), Kenneth Rainin Foundation (K.C.), Judith & Stewart Colton Center of Autoimmunity (K.C.), and the Heinz Endowments (T.S.D.). K.C. is a Burroughs Wellcome Fund Investigator in the Pathogenesis of Infectious Diseases.

## AUTHOR CONTRIBUTIONS

S.D. and K.C. conceived the study and designed the experiments. S.D. performed, analyzed, and interpreted all the experiments. T.H. and A.R.V. helped perform the viral colonization experiments. J.A.N. helped design and interpret data regarding MNV. S.Y.W. prepared the minimal defined flora. J.J.B., K.U., and T.S.D. provided the T1L virus and helped design T1L detection method. K.C. oversaw analysis and interpretation of all experiments described. S.D. and K.C. wrote the manuscript with inputs from all the authors.

## DECLARATION OF INTERESTS

K.C. receives research funding from Pfizer and Abbvie. K.C. has consulted for or received an honorarium from Puretech Health, Genentech, and Abbvie. K.C. has provisional patents, U.S. Patent Application. No. 15/625,934 and 62/935,035.

## Material and methods

### Mice

GF C57BL/6J were bred in flexible-film isolators at the New York University Grossman School of Medicine Gnotobiotics Animal Facility. Absence of fecal bacteria was confirmed monthly by evaluating the presence of 16S DNA in stool samples by qPCR as previously described (Kernbauer et al., 2014). For experiments, GF mice were housed in Bioexclusion cages (Tecniplast) with access to sterile food and water. Conventional C57BL/6J and *Rag1*^−/−^ mice were purchased from The Jackson Laboratory (Bar Harbor, ME, USA). Experiments depicted in Fig. 1 were performed using GF mice from both sexes and conventional male mice. Experiments depicted in Fig. 2-5 were performed using GF female mice. Each independent experiment comprised 8-12 mice and untreated GF mice were included in each round. Each microbial association was evaluated in 5-7 mice from at least 2 independent experiments. Littermates were randomly assigned to the experimental groups and mice were never single-housed. All animal studies were performed according to protocols approved by the NYU Grossman School of Medicine Institutional Animal Care and Use Committee.

### Virus production

MNV strains CR6 and CW3 stocks were prepared by transfecting 293T cells (ATCC) with plasmids containing the viral genome (described in (Sutherland et al., 2018)) using X-tremeGENE™ HP DNA Transfection Reagent (Roche, Indianapolis, IN, USA). Supernatants were applied to RAW264.7 cells (ATCC) for two rounds of amplification, followed by ultracentrifugation of the supernatant and resuspension in endotoxin-free PBS (Corning, Corning, NY, USA) to generate viral stocks. Concentration of stock was determined by plaque assay (described below) on RAW264.7 cells overlaid with DMEM (Corning) + 1% methylcellulose (Sigma-Aldrich, St. Louis, MO, USA) and evaluated 3 days later using crystal violet.

CVB3 strain H3 stock was prepared by transfecting HeLa cells (ATCC) with plasmids containing the viral genome and the T7 polymerase, a gift from Dr. Pfeiffer J (UT Southwestern, Dallas, TX, USA), using Lipofectamine 3000 (Thermo Fisher Scientific, Rochester, NY, USA). Cell lysate was applied to HeLa cells for two rounds of amplification. Then, the cell lysate was resuspended in PBS + 1 mM MgCl2 and 1 mM CaCl2, and freeze/thawed, and the supernatant was collected and used as viral stock. Stock titer was determined by plaque assay (described below) on HeLa cells overlaid with MEM (Lonza, Walkersville, MD, USA) + 0.5% agarose (Thermo Fisher Scientific, Waltham, MA, USA) and evaluated 3 days later using crystal violet.

MAdV1, MAdV2, and CMT93 cells were a gift from Dr. Smith JG (University of Washington, Seattle, WA, USA). Viruses were expanded on CMT93 cells and supernatants were collected and used as viral stocks. Concentration of stocks were determined by focus forming assay (described below) on CMT93 cells.

MuAstV-NYU1 stock was generated from the stool of Rag1^−/−^ mice bred at NYU Grossman School of Medicine. Briefly, stools from 6-10 weeks old mice were harvested and homogenized in PBS. Fecal slurry was pelleted, and supernatant was filtered twice using 0.22 μm Millex-GP syringe-driven filter unit (MilliporeSigma, Burligton, MA, USA). Viral titer was determined by qPCR after RNA extraction and retrotranscription.

MVMi and MVMp were a gift from Dr. Pintel D (University of Missouri, Columbia, MO, USA), and NB324K cells were a gift from Dr. Tattersall P (Yale University, New Haven, CT, USA). Viruses were expanded on NB324K cells and either cell lysate (MVMp) or supernatant (MVMi) were used as viral stocks. Concentration of stocks were determined by focus forming assay (described below) on NB324K cells.

RRV and MA-104 cells were a gift from Dr. Greenberg HB (Stanford University, Stanford, CA, USA). Virus was expanded on MA-104 cells and supernatant was collected and used as viral stock. Concentration of stock was determined by plaque assay (described below) on MA-104 cells overlaid with M199 (Sigma-Aldrich) + 0.5% agarose and evaluated 5 days later using neutral red. Reovirus T1L was prepared as described (Sutherland et al., 2018). T1L was quantified by plaque assay using L929 cells overlaid with DMEM containing 1% agar and evaluated 6 days later following neutral red staining (Sutherland et al., 2018).

### Viral inoculation

Viruses were administered to mice by oral gavage at about 5 weeks of age. Doses administered were 3×10^6^ PFU/mouse for MNV CR6 and CW3; 1×10^7^ PFU/mouse for CVB3; 1×10^6^ FFU/mouse for MAdV1; 5×10^4^ FFU/mouse for MAdV2; 1×10^10^ genome copies/mouse for MuAstV; 2×10^5^ FFU/mouse for MVMi; 5×10^6^ FFU/mouse for MVMp; 2×10^7^ PFU/mouse for RRV; 1×10^8^ PFU/mouse for T1L. For experiments depicted in Fig. 1 stool and blood were collected before viral inoculation and at 2, 5, 10, 30 and 60 days after inoculation. For experiments depicted in Fig. 2-5, stool was collected before viral inoculation and at 5 and 28 days after inoculation, whereas blood was collected 28 days after inoculation.

### Sample processing and nucleic acid extraction

Stool samples were homogenized in PBS for nucleic acid extraction by mechanical disruption with zircon beads (BioSpec Products, Bartlesville, OK, USA) using a FastPrep-24 machine (MP Biomedicals, Solon, OH, USA). Lysate slurry was spun down at 2000 g, 5 min, 4°C and the supernatant was spun down again at 8000 g, 5 min, 4°C to completely remove debris. Colon and small intestine segments were mechanically disrupted in PBS with metal beads (Qiagen) using a FastPrep-24 machine. Subsequently, lysate slurry was spun down at 8000 g, 5 min, 4°C to remove debris and a portion of the supernatant was used for RNA extraction. DNA was purified using DNeasy Blood & Tissue Kits (Qiagen) according to the manufacturer’s protocol. RNA was purified using RNeasy extraction kits (Qiagen) with a DNase (Qiagen) incubation step according to the manufacturer’s protocol. 200 μL of stool PBS homogenate and 50 μL of blood were used for nucleic acid extraction. cDNA was synthesized using ProtoScript First Strand cDNA Synthesis Kit (NEB) using random primers according to the manufacturer’s protocol. All cDNA products were stored at −20 °C.

### Viral quantification

For plaque assays, samples were serially diluted in PBS or DMEM and 500 μL were used to overlay almost confluent cells in 6 well plates (Corning). Cells were incubated at 37°C and gently shaken every 15 minutes. After 1 h, inoculum was removed, and cells were overlaid with the semi-solid media described above. After the number of days indicated above, cells were either fixed by adding PBS + 4% PFA (Sigma-Aldrich) for 1 h and then stained with crystal violet or incubated ON with PBS + 0.05% Neutral Red (Sigma, St Louis, MO, USA) and fixed with PBS + 4% PFA for 1 h.

For focus forming assays, samples were serially diluted in PBS or DMEM and 50 μL were used to overlay almost confluent cells in a 96 multi-well clear-bottom black plate (Corning). Cells were incubated at 37°C and gently shaken every 15 minutes. After 1 h, inoculum was removed and cells were overlaid with DMEM + 10% fetal bovine serum (GE Healthcare Life Science, Piscataway, NJ, USA). After 1 day, cells were fixed with HyClone Water (GE Healthcare Life Science) + 2% PFA for 20 min on ice, and then permeabilized with a quench/perm buffer (20 mM glycine, 0.25% TX-100 in PBS) for 20 min on ice. Cells were then stained with a primary antibody for 1 h on ice. Anti-adenovirus antibody clone 8C4 (Fitzgerald Industries) was used to detect MAdV1 and MAdV2), and a non-commercial α-MVM NS protein antibody previously described (Yeung et al., 1991, a gift from Dr. Tattersall P) was used to detect MVM. Then, we stained for 15 min on ice with a secondary α-mouse IgG AF488 (Thermo Fisher Scientific) and DAPI (Sigma-Aldrich). Plates were imaged using an EVOS Cell Imaging System (Thermo Fisher Scientific) and focus forming units were manually enumerated using ImageJ (NIH).

Quantification of viral nucleic acid was performed on DNA and cDNA samples using LightCycler 480 SYBR Green I Master or LightCycler 480 Probes Master (Roche), and absolute amount was calculated by comparison with in-house linearized plasmid standards. Primer and probe sequences are reported in Table S5.

### Plaque reduction neutralization test

Serum was recovered from blood collected from the submandibular vein at 20-30 days after viral inoculation. Serum inactivated for 30 min at 56°C was diluted in PBS and the same amount of virus was added to all conditions before 1 h incubation at 37°C. Then, this mix was used as inoculum for plaque assay and focus forming assay, which were performed as previously described.

### Organ processing

Colon, small intestine, mesenteric lymph nodes, lungs, and spleen were harvested from untreated GF mice or GF mice 28 days after inoculation with viruses and bacteria.

A segment of the distal colon (4 mm long and 3 cm away from the rectum) and three segments of the midsection of the duodenum, jejunum, and ileum (each 2 mm long) were collected and kept at −80°C until RNA isolation. Additionally, 3 mm from the distal colon and from the ileum were collected and fixed in formalin (Thermo Fisher Scientific) for histological analysis.

For single cells suspension, small intestinal and colonic tissues were flushed with PBS, fat and Peyer’s patches were removed, and the tissues were incubated first with 20 mL of HBSS (Gibco) with 2% HEPES (Corning), 1% sodium pyruvate (Corning), 5mM EDTA, and 1 mM dithiothreitol (Sigma-Aldrich) for 15 min at 37°C, and then with new 20 mL of HBSS with 2% HEPES, 1% sodium pyruvate, 5mM EDTA for 10 min at 37°C. Tissue bits were washed in HBSS + 5% FCS, minced, and then enzymatically digested with collagenase D (0.5 mg/mL, Roche) and DNAse I (0.01 mg/mL, Sigma-Aldrich) for 30-45 min at 37°C with constant stirring. Digested solutions were passed through 70 μm cell strainers (BD) and cells were subjected to gradient centrifugation using 40% Percoll (Sigma-Aldrich).

IELs were recovered from the liquid phase of the first small intestine incubation, washed with PBS, and subjected to gradient centrifugation using 40% Percoll.

mLNs were collected and passed through 100 μm cell strainers and resuspended in PBS.

Lungs and spleens were grossly minced and enzymatically digested with collagenase D (0.5 mg/mL) and DNAse I (0.01 mg/mL) for 20-30 min at 37°C. Digested solutions were passed through 100 μm cell strainers, resuspended in ACK buffer to lyse the red blood cells, and resuspended in PBS.

For the analysis of the cytokine production, cells were plated in RPMI with 10% FBS and treated with phorbol 12-myristate 13-acetate (50 ng/mL, MilliporeSigma) and ionomycin (1 μg/mL, MilliporeSigma) in the presence of *GolgiStop* (BD) and *GolgiPlug* (BD) for 4 h at 37°C.

### Flow Cytometry

Cells were pre-incubated with CD16/CD32 Fc block (BD PharMingen). Surface and intracellular cytokine staining was performed per manufacturer’s instructions in PBS + 2% FBS for 20 min on ice. Three staining panels were utilized. The first panel included antibodies against BST2, NK1.1, THY1.2, F4/80, CD103, LY6C, CD11b, MHC-II, CD45, CD11c, CD19, CD64, and B220. To stain the spleen samples, we substituted CD103 with CD8a for a better evaluation of the dendritic cell subsets. The second panel included antibodies against GATA3, CD11b, CD11c, GR1, CD19, TER119, Tbet, TCRγδ, FOXP3, CD8, CD4, RORγt, CD62L, CD127, NK1.1, CD44, CD3ε, CD45. The third panel included antibodies against IFN-γ, CD11b,CD11c, GR1, CD19, TER119, Nk1.1, Il-22, TCRγδ, GRANZYME B, IL-17a, CD8, CD4, IL-10, CD127, IL-4, CD3Ε, CD45. Samples were fixed with either Fixation Buffer (Biolegend, San Diego, CA, USA) or eBioscience Foxp3/Transcription Factor Staining Buffer Set (Thermo Fisher Scientific). For intracellular staining of transcription factor, cells were permeabilized with the eBioscience Foxp3/Transcription Factor Staining Buffer Set at room temperature for 30 min in the presence of antibodies. For intracellular staining of cytokines, cells were permeabilized with Intracellular Staining Permeabilization Wash Buffer (Biolegend) at room temperature for 30 min in the presence of antibodies. Zombie UV Fixable Viability Kit (Biolegend) was used to exclude dead cells. Samples were acquired on a BD LSR II (BD Biosciences) and analyzed using FlowJo software (Treestar, Inc., Ashland, OR, USA).

### RNA deep sequencing

CEL-seq2 was performed on 67 colonic and 60 small intestinal RNA samples. Sequencing was performed on Illumina NovaSeq 6000 (Illumina). All samples from the same organs were sequenced together, thus no correction for batch effect was necessary.

### Microscopy on intestinal tissue

Small intestinal and colonic tissues were cut open along the length, pinned on black wax, and fixed in 10% formalin. Tissues were embedded in 3% low melting point agar (Promega, Madison, WI, USA). Formalin embedding, cutting, and hematoxylin and eosin staining was performed by the NYU Histopathology core. Sections were imaged either on a Leica SCN400 F microscope (Leica Biosystems, Buffalo Grove, IL, USA).

### Bacteria

The Minimal Defined Flora consisted of the 15 bacteria described in Brugiroux et al., 2016. *Akkermansia muciniphila* YL44 was a gift from Dr. McCoy K (University of Calgary, Canada) and it was grown in 0.1% mucin (Sigma-Aldrich), anaerobic, 37°C. *Bacteroides caecimuris* I48 was from DSMZ and it was grown in BHI (Anaerobe Systems,), anaerobic, 37°C. *Muribaculum intestinale* YL27 (DSMZ) was grown in chopped meat media (Anaerobe Systems), anaerobic, 37°C. *Turicimonas muris* was a gift from Dr. McCoy K and it was grown in BHI, anaerobic, 37°C. *Escherichia coli* Mt1B1 (DSMZ) was grown in LB (Sigma-Aldrich), aerobic, 37°C. *Bifidobacterium longum* subsp. animalis YL2 (DSMZ) was grown in BHI, anaerobic, 37°C. *Staphylococcus xylosus* 33ERD13C (DSMZ) was grown in TSB-yeast (Sigma Aldrich), aerobic, 37°C. *Streptococcus danieliae* ERD01G (DSMZ) was grown in TSB-yeast, microaerophilic, 37°C. *Enterococcus faecalis* KB1 (DMSZ) was grown in TSB-yeast, aerobic, 30°C. *Acutalibacter muris* KB18 (DSMZ) was grown in BHI, anaerobic, 37°C. *Clostridium clostridioforme* YL32 (DSMZ) was grown in PYG (Anaerobe Systems), anaerobic, 37°C. *Flavinofractor plautii* YL31 (DSMZ) was grown in PYG, anaerobic, 37°C. *Blautia coccoides* YL58 (DSMZ) was grown in chopped meat media, anaerobic, 37°C. *Lactobacillus reuteri* I49 (DMSZ) was grown in MRS, microaerophilic, 37°C. *Clostridium innocuum* I46 (DSMZ) was grown in chopped meat media or PYG, anaerobic, 37°C.

*Yersinia Pseudotubercolosis* was a gift from Dr. Darwin A (NYU), and it was grown overnight in Luria-Bertani broth with shaking at 28°C. In the morning, the bacterial were subcultured in fresh Luria-Bertani broth with shaking at 28°C until OD 0.7-0.9. Bacterial density was confirmed by dilution plating. 9-week-old female GF mice were inoculated by oral gavage with 2×10^4^ CFU resuspended in 200 μl PBS. Severity of disease was quantified through a scoring system in which individual mice received a score of 1 in case of the presence of visible blood in the stool, and between 0 and 2 of the following: hunched posture and diarrhea.

## Quantification and statistical analysis

### Immunophenotypes

Flow cytometry fold change values were calculated by dividing the frequency of a given cell type by the average frequency obtained from the GF mice in the same experimental round. Statistical differences between each colonization condition and the GF mice were calculated by one-way ANOVA followed by Dunn’s post-hoc analysis using the R package “stats”. To control for multiple testing, a false discovery rate was calculated by the Benjamini-Hochberg procedure using the R package “stats” for each cell type analyzed.

### Selection of differentially expressed genes

RNA-Seq results were processed using the R package “DESeq2” to obtain variance stabilized count reads, fold changes relative to GF condition, and statistical p-value. Analysis of the whole tissue transcriptome focused on differentially expressed genes, defined as the genes with an absolute fold change relative to GF >2 and an unadjusted p-value <0.01.

### Computational analysis

Hierarchical clustering of the population and cytokine frequencies were performed on the Euclidean distances using the R package “stats”. Distance-based redundancy analysis (db-RDA) was used to determine the contribution of different factors to the variance observed within the immunophenotypes samples or differentially expressed genes using the R package “vegan”. Euclidean distance between colonization conditions according to differentially expressed genes was calculated using the R package “stats”, and permutational multivariate analysis of variance on these distances was calculated using the R package “vegan”. Heatmaps were generated using either the package “ggplot2” or “pheatmap”. Gene ontology analysis was performed using the package “clusterProfiler”. GSEA was performed using the package “WebGestaltR”. Canonical pathway and upstream regulators analysis were performed by uploading the differentially expressed genes to Ingenuity Pathway Analysis software (Qiagen).

### GSEA gene signatures

GSEA gene signatures were generated in a manner similar to Godec et al., 2016 by selecting the top upregulated or downregulated genes, up to 200, with an FDR<0.02 or an unadjusted p-value<0.001. Gene signatures consisting of less than 10 genes were discarded. IL-22 and bacterial signatures were based on the transcriptional data described in Gronke et al., 2019 and Geva-Zatorsky et al., 2017, respectively.

### Statistical analysis

Statistical differences were determined as described in figure legend using either R or GraphPad Prism 8 software (La Jolla, CA, USA).

## Data and software availability

The extensive datasets presented in this manuscript are made available in Tables S1-S4. The immunophenotypes are presented in Table S1C as frequencies of cell types and in Table S1B as` fold changes relative to uninfected GF mice. The accession number for the gene expression raw data reported in this paper is pending.

---

**Figure S1.**
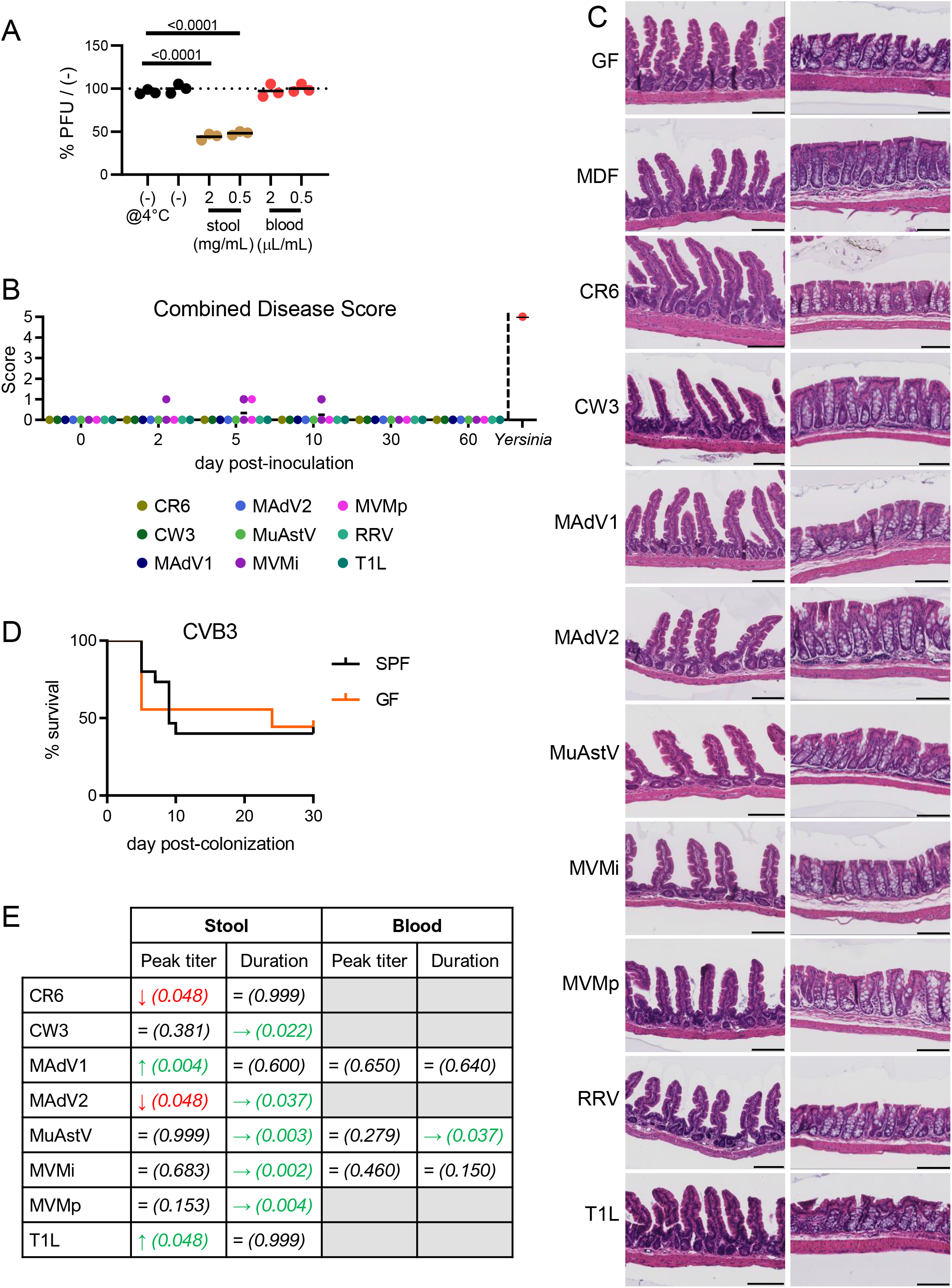
Enteric Virus Infection in Conventional and GF Mice. (A) Incubation of RRV at 37°C for 1 h with clarified stool lysate but not blood lead to a reduction in plaque forming units (PFUs). Values were normalized to the number of plaques observed when the same stock of RRV was kept at 4°C in parallel. Data are representative of two independent experiments. Dots depict replicates from one representative experiment. Statistical significance was calculated by ANOVA followed by Dunn’s post-hoc analysis. (B) GF mice inoculated with the enteric viruses shown in Figure 1 were scored at indicated days post-inoculation (dpi) for diarrhea (0: no diarrhea, 2: watery stool), hunched posture (0: no hunching, 2: hunched), and visible blood in the stool (0: no, 1: yes). As a reference, the combined disease score is shown for four GF mice orally inoculated with *Yersinia Pseudotubercolosis* on day 7 from two independent experiments. (C) H&E-stained sections of the small intestine (left) and colon (right) of GF mice at 28 dpi indicating absence of overt inflammation. Bar indicates 100 μm. (D) Survival of 15 conventional and 9 GF mice inoculated perorally with CVB3 from four independent experiments. (E) Time course of viral loads in stool and blood of conventional versus GF mice from Figure 1 were compared to identify significant differences in peak titer and duration. Statistical significance for peak titer was calculated using a non-parametric Mann-Whitney test at the timepoint with the highest viral titer in GF mice. Green upward arrow and red downward arrow refer to an increase and decrease in viral titer in GF mice, respectively. Statistical significance for the duration of viral shedding was calculated using a log-rank test. Green right-facing arrow refers to prolonged detection of virus in GF mice. Gray boxes indicate conditions in which viruses were not detected in the blood. Value in parentheses denote p-values.

**Figure S2.**
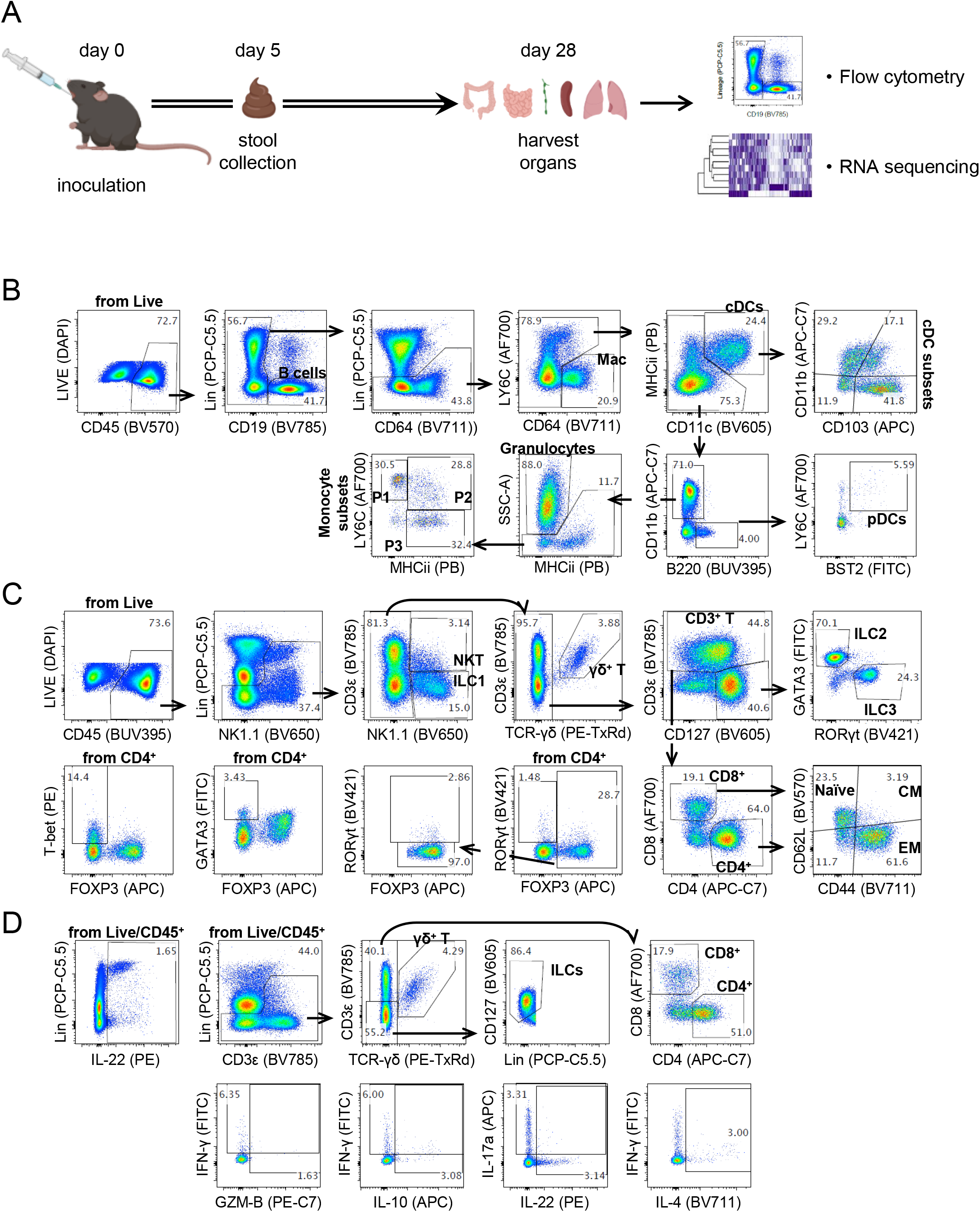
Experimental Design and Gating Strategy. (A) Experimental design for immune profiling of virus-infected mice. Five-to-six-week-old mice were untreated or inoculated with viruses or bacteria. Four weeks after inoculation, effects on immune cells and the transcriptome was evaluated. Five-to-eight mice were analyzed per condition. To ensure that each condition was represented in at least two independent experiments, results of 11 independent experiments with 8-12 mice are presented and include control untreated GF mice in each independent experiment. Prepared using BioRender.com. (B-D) Flow cytometry gating strategies for B cells and myeloid cells (B), T cells and ILCs (C, transcription factors), and cytokine production (D).

**Figure S3.**
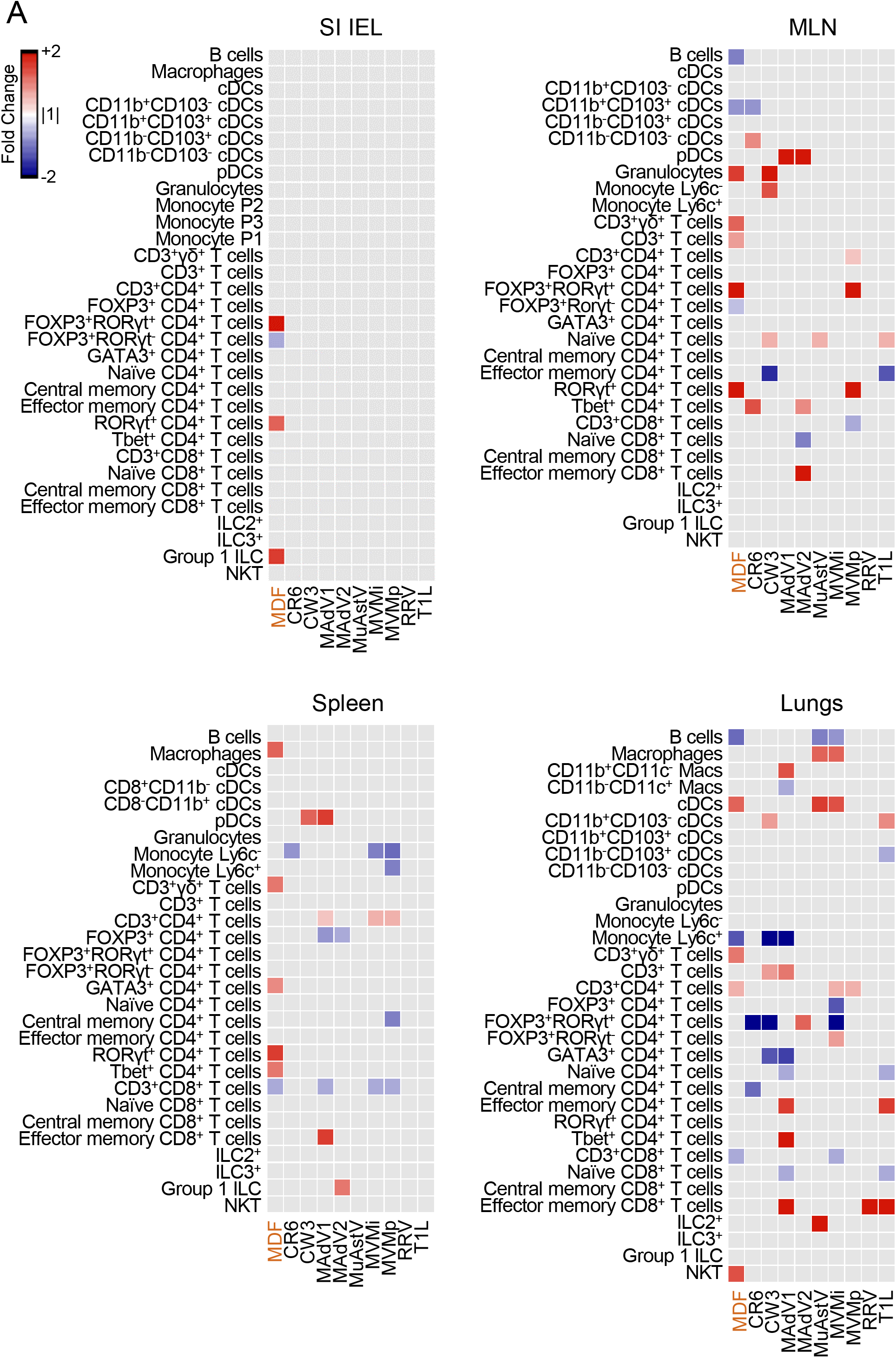
Enteric Viruses Promote Changes in Immune Cell Populations of Extra-Intestinal Tissues. Heatmap showing the average fold-change for each immune population relative to GF in IELs, mLNs, spleen, and lungs with an FDR<0.1. Gray: FDR>0.1.

**Figure S4.**
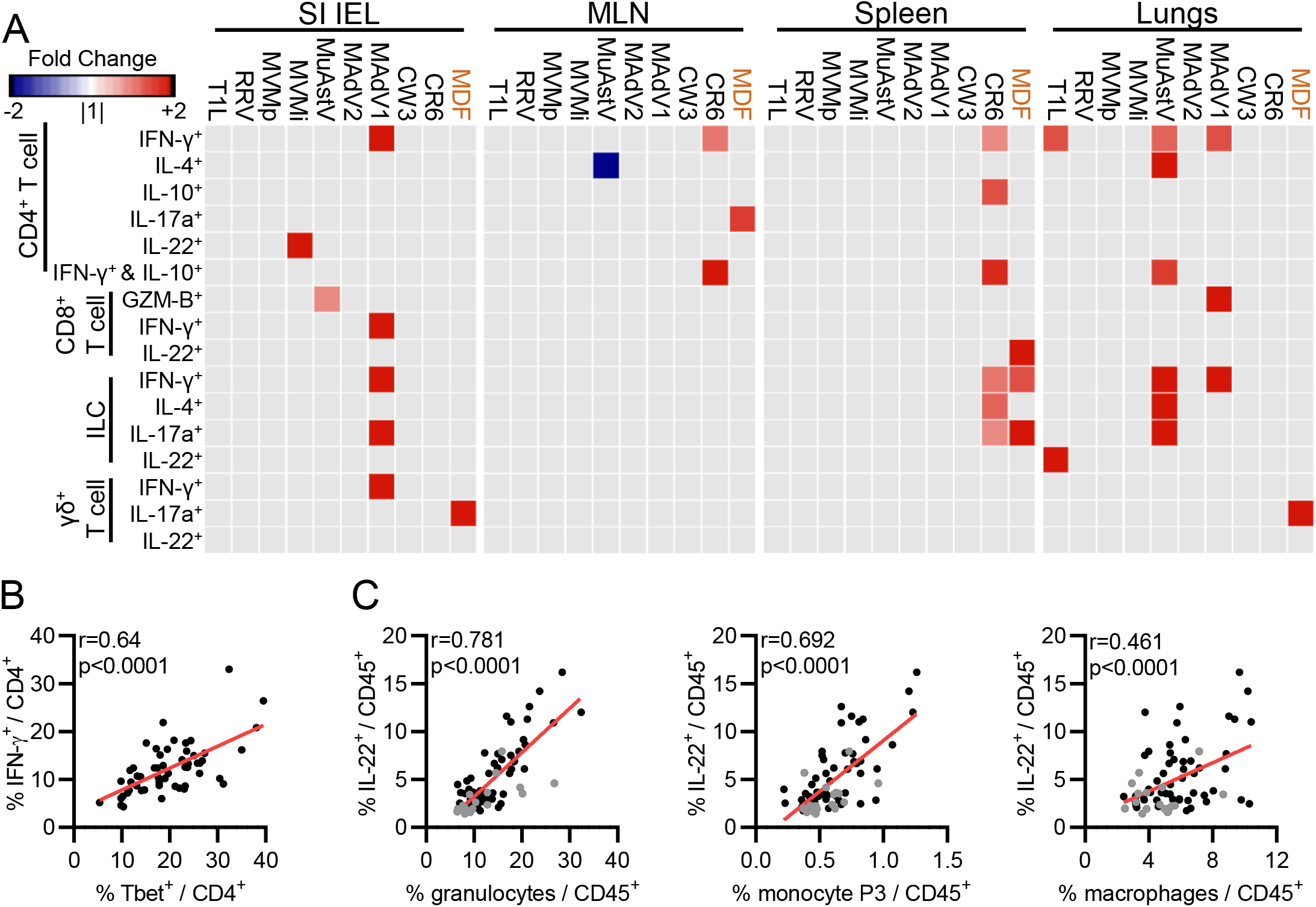
Enteric Viruses Increase Cytokine Production by Immune Cells in Extra-Intestinal Tissues. (A) Heatmaps showing average fold-change for cytokine-producing immune cell populations for the indicated conditions relative to GF mice in IELs, mLNs, spleen, and lungs with an FDR < 0.1. Gray: FDR > 0.1. (B-C) Pearson correlation between the indicated population frequencies in cLP. Black dots: virus-infected samples; gray dots: GF samples.

**Figure S5.**
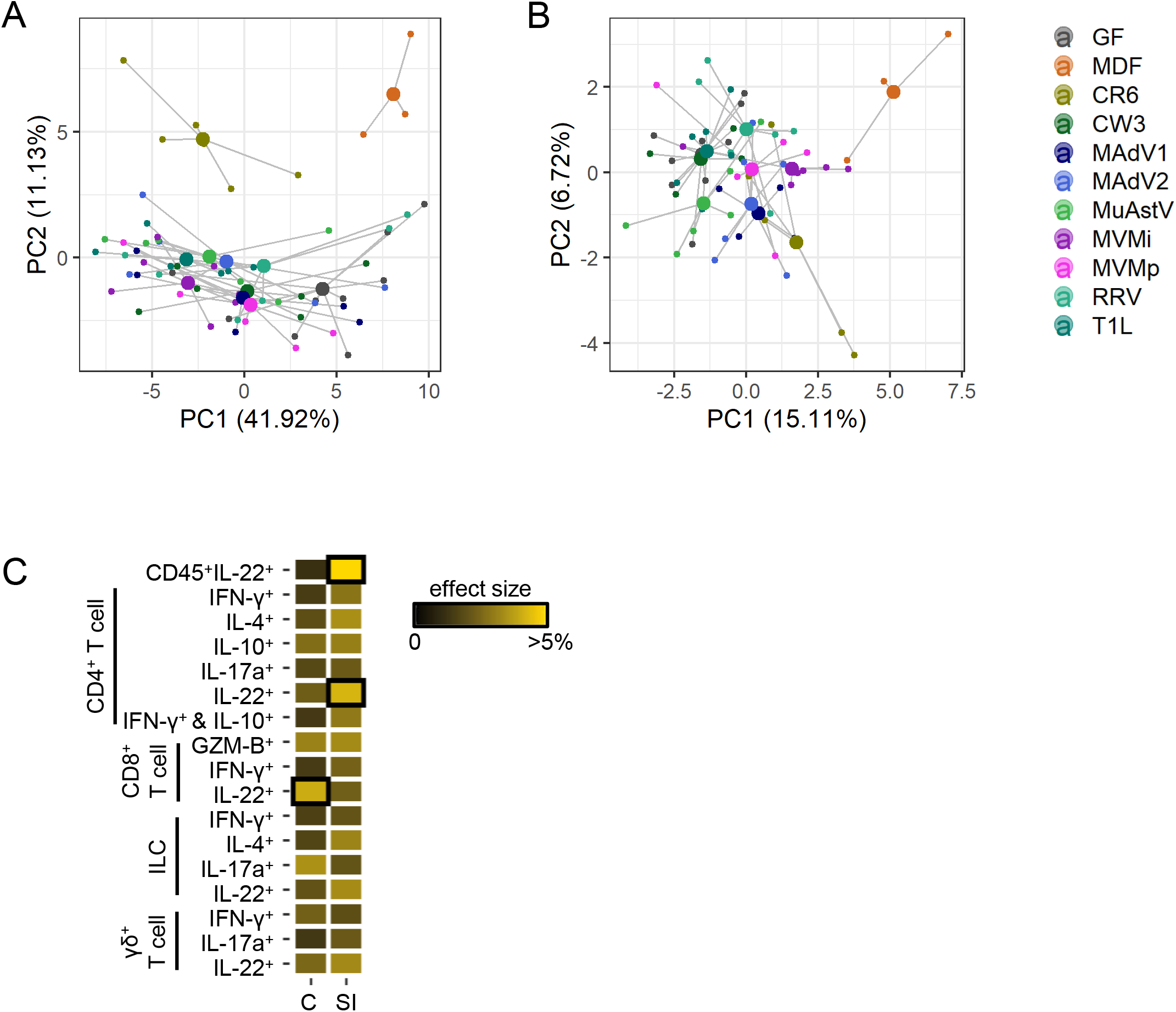
Tissue Transcriptome Induced by Viral Infection. (A-B) PCA clustering of the colonic (A) and small intestinal (B) DE genes. Samples inoculated with the same microbe are connected by lines to the calculated group centroids. (C) db-RDA indicating the individual effect size of the cytokine production frequencies obtained by flow cytometry as explanatory variables of the DE gene variance in the colon and small intestine. Black boxes indicate p<0.05.

**Figure S6.**
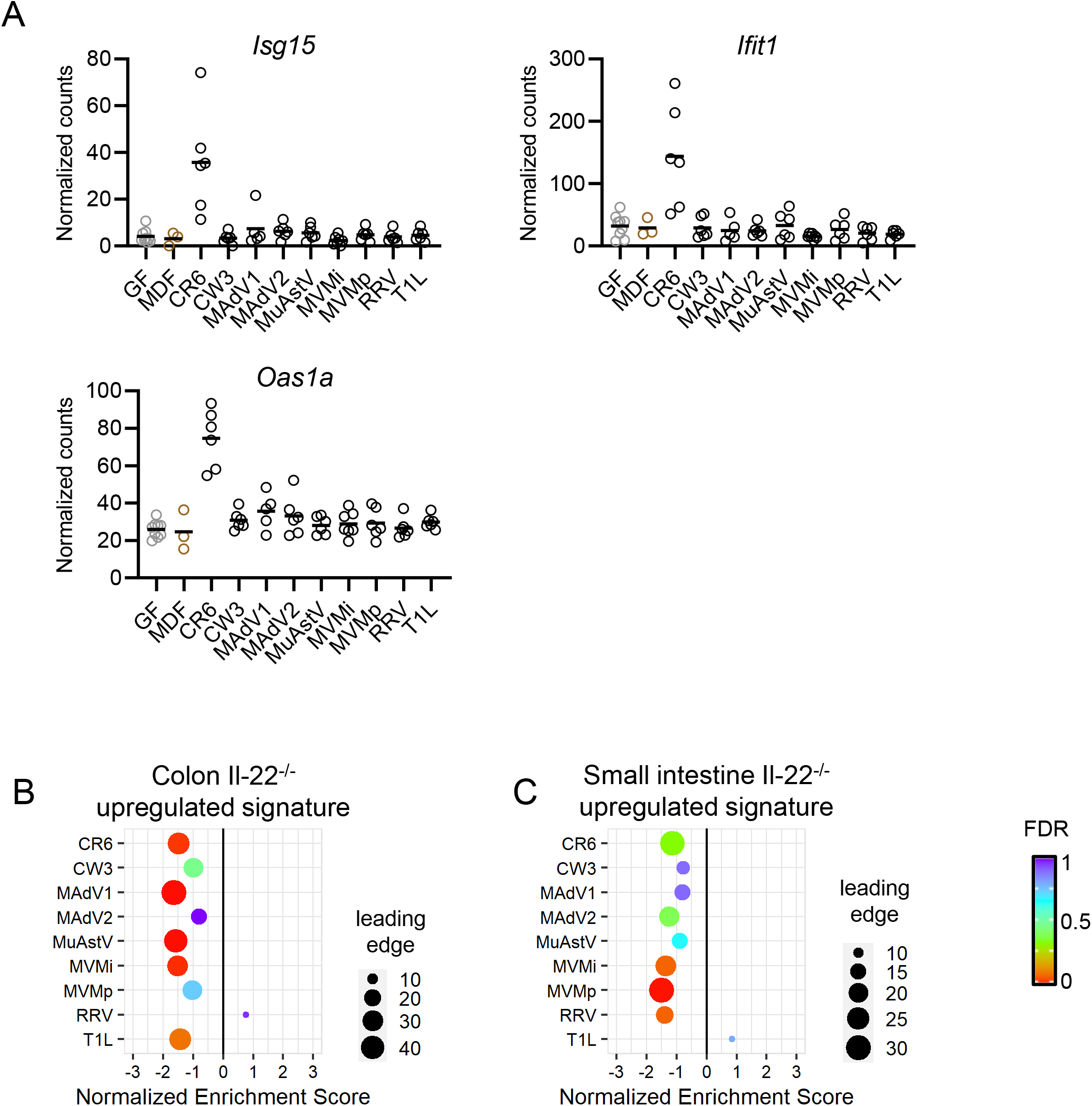
Expression of Interferon-Stimulated Genes and IL-22 Signature. (A) DESeq2 transformed counts of the indicated representative interferon-stimulated genes (ISGs) from RNA-Seq of the colon. (B-C) Gene expression in the colon (B) and small intestine (C) from virus-infected mice were analyzed for enrichment of transcripts upregulated in the intestines of IL-22−/− mice by GSEA.

**Table S5:**
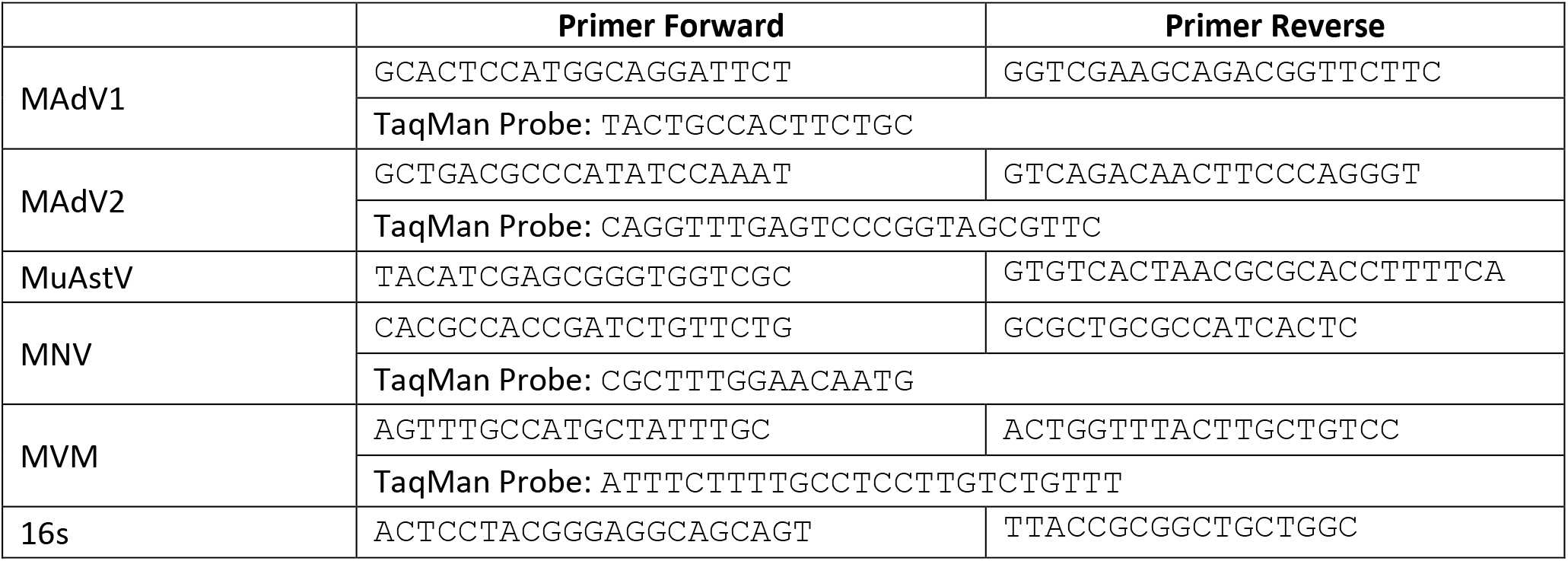
Primers and probes used in the study.

**Table S6:**
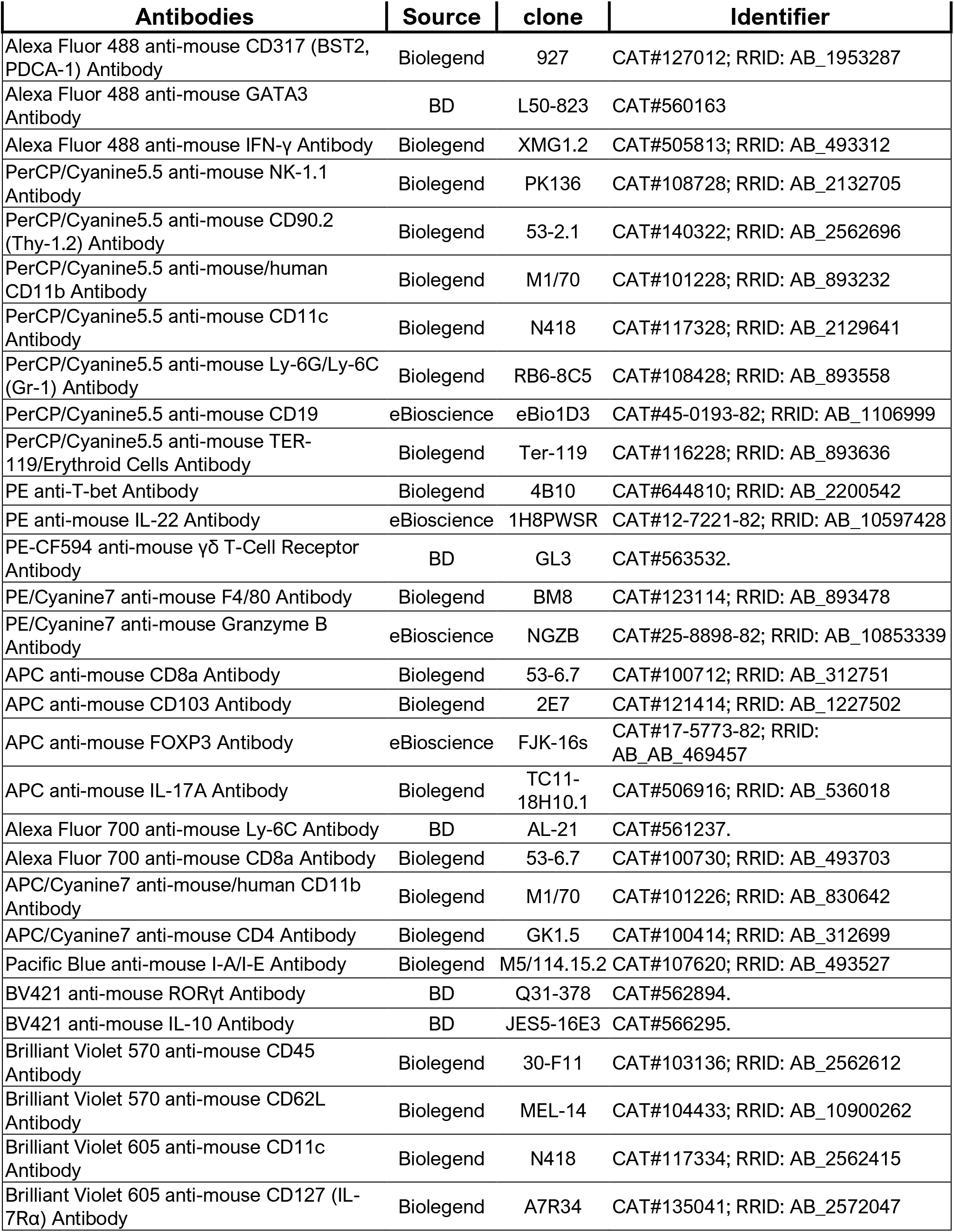
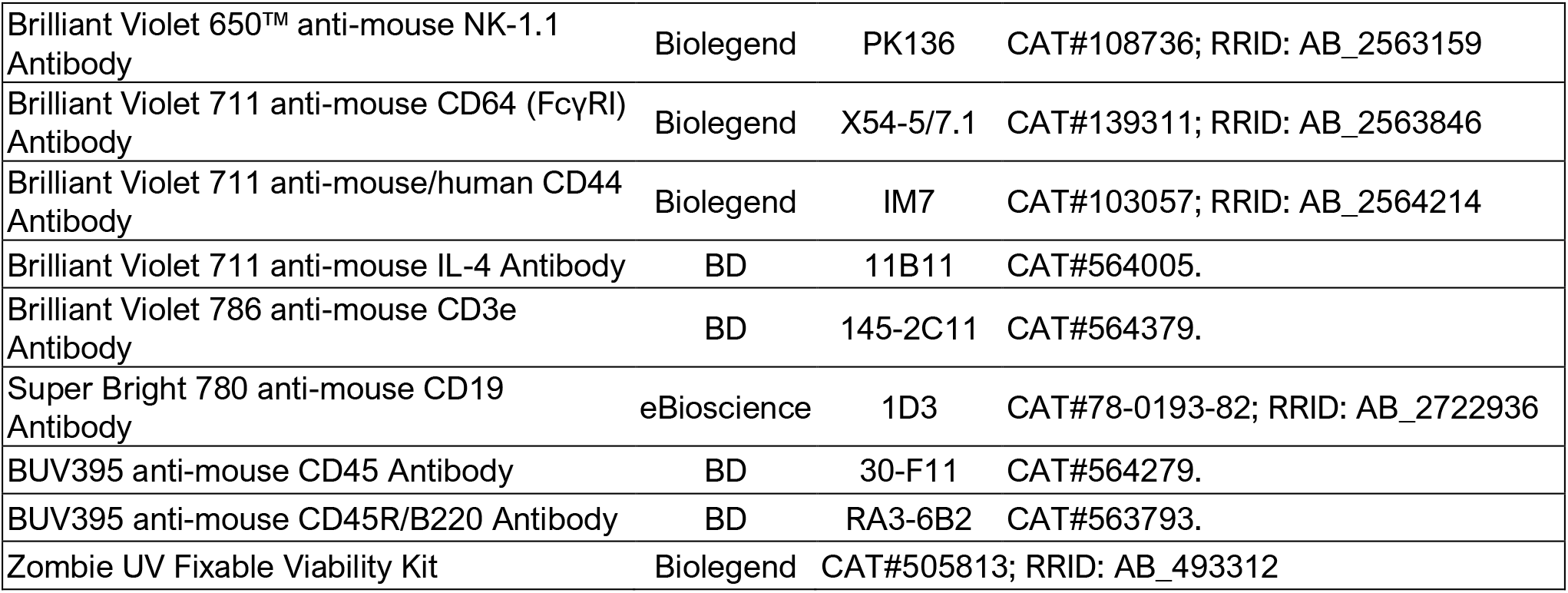
List of the flow cytometry antibodies used in this study.

